# Reprogramming SREBP1-dependent lipogenesis and inflammation in high-risk breast with licochalcone A: a novel path to cancer prevention

**DOI:** 10.1101/2024.05.20.595011

**Authors:** Atieh Hajirahimkhan, Elizabeth T. Bartom, Carolina H Chung, Xingyu Guo, Kyli Berkley, Oukseub Lee, Ruohui Chen, Wonhwa Cho, Sriram Chandrasekaran, Susan E. Clare, Seema A. Khan

## Abstract

**Background:** Anti-estrogens have had limited impact on breast cancer (BC) prevention. Novel agents with better tolerability, and efficacy beyond estrogen receptor (ER) positive BC are needed. We studied licochalcone A (LicA) for ER-agnostic BC prevention.

**Methods:** We evaluated antiproliferative effects of LicA in seven breast cell lines and its suppression of ER+ and ER− xenograft tumors in mice. High-risk human breast tissue was treated with LicA *ex vivo*, followed by RNA sequencing and metabolism flux modeling. Confirmatory testing was performed in an independent specimen set and ER+/− BC cell lines using NanoString metabolic panel, proteomics, western blots, and spatiotemporally resolved cholesterol quantification in single cells.

**Results:** LicA suppressed proliferation *in vitro* and xenograft tumor growth *in vivo*. It downregulated pivotal steps in PI3K-AKT-SREBP1-dependent lipogenesis, suppressed PI3K and AKT phosphorylation, SREBP1 protein expression, and cholesterol levels in the plasma membrane inner leaflet, to the levels in normal breast cells. LicA also suppressed prostaglandin E2 synthesis and PRPS1-catalyzed *de novo* nucleotide biosynthesis, stalling proliferation; further evident by reduced MKI67 and BCL2 proteins.

**Conclusions:** LicA targets SREBP1, a central regulator of lipogenesis and immune response, reducing pro-tumorigenic aberrations in lipid homeostasis and inflammation. It is a promising non-endocrine candidate for BC prevention.

## Background

Current recommendations for BC prevention are based on estimates of risk in individual women [1]; for those at markedly increased risk due to high-penetrance germline mutations, risk-reducing bilateral mastectomy can be considered. However, the vast majority of at-risk women are at risk for other reasons, with risk estimates in the 1.5-5-fold range. For them, risk-reducing medications are recommended. At present, the only risk-reducing drugs available for BC are anti-estrogens; aromatase inhibitors (AIs) for postmenopausal women and selective estrogen receptor modulators (SERMs) for pre and postmenopausal women. These drugs can lower the risk of estrogen receptor positive (ER+) BC by 50-65% [2]. However, of the estimated 10 million U.S. women who could benefit from these medications more than 85% decline them, mainly due to their adverse effects: vasomotor symptoms, thrombogenesis/stroke, bone loss, and endometrial cancer [2–7]. In addition, these risk reducing drugs do not prevent estrogen receptor negative (ER-) BC. One of the pathways abnormally active in precancerous lesions such as atypical hyperplasia and ductal carcinoma in situ (DCIS) is PI3K-AKT [8–10]. PI3K pathway inhibitors such as everolimus and alpelisib have been proposed for repurposing as ER− and ER+ BC prevention agents [11–13]; however, their side effects such as hyperglycemia are a major impediment [14]. There have been attempts to repurpose widely used and well-tolerated drugs such as metformin for BC risk reduction; however, clinical trial results have been disappointing [15, 16]. Therefore, there is a clear need to have alternative agents with sufficient efficacy, minimal adverse effects, and greater acceptance.

Upregulated *de novo* fatty acid and cholesterol biosynthesis mediated by the activation of the PI3K-AKT-SREBP axis has been shown to have an important role in BC development and progression through enhanced proliferation, inter and intracellular signaling, and enabling evasion from immune surveillance [17, 18]. Correspondingly, metabolic dysregulation and obesity are associated with the elevated risk: ER+ disease in post-menopausal women and ER− BC in those who are premenopausal [19–24].

The activity of the master regulator of lipogenesis, sterol regulatory element binding protein 1 (SREBP1) has been shown to have prognostic value in BC progression [25, 26]. SREBP1 plays a major role in macrophage polarization and regulation of inflammasome in the tissue microenvironment and forms a complex with NF-kB that regulates the inflammatory response [27, 28]. Excess activation of SREBP1 upregulates the expression of most fatty acid biosynthesis enzymes and SREBP2-dependent cholesterol biosynthesis [29]. This subsequently promotes an inflammatory/oxidative microenvironment, provides the excess cell membrane structural components necessary for proliferating cells, protects malignant cells from immune destruction, promotes cancer stem cell viability, and supplements cholesterol and its derivatives as precursors for estrogen biosynthesis [30–33]. In addition, the role of dysregulated lipogenesis in promoting ER− BC is associated with the inflammatory cytokines secreted from immune cells recruited to the stroma consequent to excess free fatty acids and cholesterol in the microenvironment, and the dysregulation of metabolic hormones and growth factors [33–35]. The cancer-promoting effects of inflammation [36–38] can be tempered through downregulating SREBP1 and through the activation of NRF2 and suppression of NF-kB-dependent pathways. [28, 39]. This presents an opportunity for interventions that prevent malignant transformation in the breast. Importantly, such a strategy will not produce deleterious endocrine responses that are of concern with presently available risk reduction drugs [39].

Our current research reveals that licochalcone A (LicA), a compound that is already in clinical trials for the relief of inflammatory skin conditions, is an excellent candidate as a non-endocrine BC risk-reducing drug with minimal toxicity and better acceptance. It has antioxidant and anti-inflammatory effects [40–52], moderately inhibits aromatase (CYP19A1), the rate-limiting enzyme in estrogen biosynthesis, and the target of AIs used for BC prevention [41]. In addition, LicA blocks estrogen carcinogenic metabolism through the downregulation of CYP1B1, *in vivo* [45]. LicA has also been reported to relieve conditions such as arthritis, mastitis, and acute lung injury through anti-inflammatory pathways, and appears to have osteoprotective effects [48, 49, 52–54]. The NF-kB-dependent anti-inflammatory effects of LicA can be partly explained by its binding to cysteine residues in KEAP1 which simultaneously enhances NRF2-dependent antioxidant effects [42] while suppressing inflammation [55], but it can also be a consequence of its effects on SREBP1-dependent pathways [28]. In the current study, we confirm our previous *in vitro* and *in vivo* antioxidant observations using LicA in breast microstructures from women at high risk of BC. We also present novel findings regarding its effects on metabolic pathways involving PI3K-AKT-SREBP1 which we have elucidated in various breast cell lines, human *ex vivo*, and *in vivo* models using multiple orthogonal approaches. In addition, we show its antiproliferative effects resulting from changes in metabolic flux, particularly through decreasing the *de novo* nucleotide biosynthesis step of the pentose phosphate pathway (PPP). Based on these results and existing literature, we posit that reducing lipogenesis and inflammation through suppressing SREBP1 can reduce the risk of BC and that LicA presents as an excellent candidate for BC prevention and is highly likely to be safe and well-tolerated.

## Methods

### Chemicals and Materials

All chemicals and reagents were purchased from Sigma-Aldrich (St. Louis, MO), unless otherwise indicated. MammoCult media kit, heparin, and hydrocortisone were purchased from Stem Cell Technologies (Vancouver, BC. Canada). F12-K nutrient mix (Kaighn’s) medium was acquired from Gibco (Dublin, Ireland). Fetal bovine serum (FBS) was purchased from Atlanta Biologicals (Norcross, GA). Collagenase I was purchased from Sigma-Aldrich (St. Louis, MO). Licochalcone A (LicA) was acquired from Med Chem Express (Monmouth Junction, NJ). Direct-Zol RNA prep kit was acquired from Zymoresearch (Irvine, CA). TRIzol was obtained from Invitrogen (Waltham, MA). RNAeasy mini prep kit was purchased from QIAGEN (Germantown, MD). RNA concentration and clean up kit was acquired from Norgen Biotek (ON, Canada). KAPA RNA HyperPrep library preparation kit was obtained from Roche (Madison, WI). Unique dual adaptors for next generation sequencing (NGS) and PCR reagents, primers, and master mix were purchased from Integrated DNA Technologies (Coralville, IA). Antibodies were obtained from Abcam (Cambridge, UK). Western blot reagents, buffers, supplies were purchased from Thermo Fisher Scientific (Hanover Park, IL.).

### Cell Culture

We grew pre-malignant DCIS.COM and malignant MCF-7 and MDA-MB-231 cells (all obtained from ATCC) in RPMI-1640 supplemented with 10% fetal bovine serum (FBS) and 1% penicillin/streptomycin (Invitrogen Thermo Fisher Scientific, Hanover Park, IL.). We also grew pre-malignant DCIS.COM/ER+ PR+ cells obtained from Dr. Dean Edwards’s laboratory (Baylor College of Medicine) [56] in DMEM/F12 supplemented with 10% horse serum, 0.5% penicillin/streptomycin, and 1% HEPES. In addition, we acquired MCF-7 cells overexpressing aromatase (MCF-7aro) from Dr. Shiuan Chen’s laboratory (City of Hope) [57] and grew them in MEM supplemented with 10% FBS, 2 mM L-glutamine, 1 mM sodium pyruvate, 1% nonessential amino acids, and 1 U/mL penicillin/streptomycin. BRCA defective malignant breast cell lines HCC-1937 and HCC-3153 were obtained from Dr. Gazdar’s group (University of Texas Southwestern Medical Center) and were grown in RPMI-1640 supplemented with 5% FBS and 1% penicillin/streptomycin.

### Human breast microstructure preparation and treatment

We prepared breast microstructures from the unaffected contralateral breast of postmenopausal women who underwent bilateral mastectomy for unilateral BC, using the protocol of Tanos et. al., with minor modifications [41, 58, 59]. The tissues were obtained while fresh and diced to approximately 5 mm pieces before digesting at 37°C with 2% collagenase I in F12-K nutrient mix (Kaighn’s) medium, overnight. After the completion of digestion, the tissue was centrifuged at 250 x g for 5 min and the supernatant was removed. The pellet was washed with phosphate buffered saline and was mixed and cultured in MammoCult media supplemented with 0.2% heparin and 0.5% hydrocortisone. After 24 h incubation at 37°C the treatments were mixed with fresh MammoCult media and added to the microstructures. **Ethics declarations**: Institutional Review Board (IRB) approval (IRB STU00202331) was obtained from Northwestern University prior to obtaining informed consents from patients and collecting samples. All experiments were conducted in accordance with the approved protocol and guidelines.

### RNA sequencing

After 24 h incubation with LicA (5 µM) or DMSO, the microstructure pellets obtained from 6 subjects were prepared and washed with HBSS. Trizol was added to each pellet and the protocol of Direct-Zol RNA prep kit was used to extract RNA. The protocol included a step of DNAse I treatment for removing DNA contamination of the RNA preparations. RNA was eluted in nuclease free water and quality assessment was performed using Agilent Bioanalyzer and Qubit. Libraries were made using Roche KAPA Biosystems protocol (KAPA RNA Hyper Prep Kit Technical Data Sheet, KR1352 – v4.17, Roche Corporate) and the quality was evaluated. Total RNA sequencing was performed with Illumina NovaSeq 6000 v1.5, 2×100 b, paired end with 40 million reading depth, followed by quality assessment using FASTQC, alignment to reference genome hg38, raw data processing using STAR, and data analysis to define differential gene expression, as described previously [60–62]. Significantly (adj P < 0.05) modulated genes were further analyzed with Enrichr to define highly modulated pathways. The sample relationships were validated with NGSCheckMate as well.

### NanoString metabolism pathway panel

We treated six additional subjects’ breast microstructures and two cell lines MDA-MB-231 (ER-) and MCF-7 (ER+) cells with LicA (5 µM and 10 µM) or DMSO for 24 h. After extracting RNA and quality assessment we used the NanoString nCounter Metabolism Panel (NanoString Technologies Inc.) to quantify RNA expression from 749 genes. Differentially expressed genes and pathway analysis were determined using ROSALIND platform (ROSALIND, Inc.) with normalization, fold changes (≥ 1.5 fold: upregulated or ≤ −1.5 fold: downregulated), and adjusted p-value < 0.05 using the Benjamini-Hochberg method.

### Genome-scale metabolic network modeling and flux-based analysis (FBA)

We calculated the relative activity of reactions in breast microstructures of 6 women by interpreting gene expression data using the Recon1 human metabolic model outlined in Shen et al [63–65]. We then identified a metabolic flux state that is most consistent with gene expression data in control and LicA treated samples. This was achieved by maximizing the activity of reactions that are associated with upregulated genes and minimizing flux through reactions that are downregulated in a condition, while simultaneously satisfying the stoichiometric and thermodynamic constraints embedded in the model using linear optimization [63–65]. The glucose, fatty acid, and glutamine levels in the simulations were adjusted based on the growth media used for culturing the high-risk women’s breast microstructures. All p-values were corrected for multiple comparisons.

### Proliferation assay

We seeded pre-malignant DCIS.COM and DCIS.COM/ER+ PR+ cell lines as well as malignant MCF-7 (ER+ PR+), MDA-MB-231 (ER− PR-), MCF-7aro (ER+ PR+), HCC-1937 (ER− PR-, BRCA1 mutated), and HCC-3153 (ER− PR-, BRCA mutated1) cells with the density of 2.5 x 10^3^ cells/well in their appropriate media in 96-well plates. We placed the plate in IncuCyte instrument for continued live cell imaging every 6 h. When cells reached 30% confluency, eight different concentrations of LicA ranging from 350 nM to 40 µM were added to the designated wells. We added comparable dilutions of DMSO to the wells allocated for vehicle control. In the case of MCF-7aro cells which overexpress *cyp19a1* (aromatase), the dosing of LicA or DMSO was performed in the presence of androstenedione, the substrate for aromatase. We continued live cell imaging for a minimum of 4 days post treatment. When we found out that a single low dose is not sufficient for sustained antiproliferation, we added the treatment and control every 48 h for 6 days and continued live cell imaging for a maximum of 12 days. Data was analyzed using Zoom software to quantify the images and were plotted as mean ± SEM of at least two independent measurements.

### Proteome Integral Solubility Alterations (PISA) proteomics

MCF-7 and MDA-MB-231 cells were treated with LicA (10 µM) for 24 h. We used the sample preparation and PISA protocol as described previously [66], with minor modifications. Cells were detached with non-enzymatic dissociation buffer and centrifuged at 300 x g for 4 min. The cell pellets were reconstituted in PBS containing protease and phosphatase inhibitors. This step was repeated. The final pellet was resuspended in PBS with inhibitors and was distributed into PCR tubes. Thermal denaturation was performed with a steady temperature at 40, 42, 45, 50, 55, 60, 63, and 65°C for 3 mins in parallel in a thermal cycler. Immediately after heating, the samples were removed and incubated at room temperature for 3 min before they were snap-frozen with liquid nitrogen. Two cycles of freeze-thaw using liquid nitrogen and a heating block set at 25°C were conducted to ensure a uniform temperature between tubes. After the vortex, the lysates were centrifuged at 20,000 x g for 20 min at 4°C to pellet the debris with precipitated and aggregated proteins. While on ice, the supernatant with the soluble protein fraction was carefully removed to a new tube. BCA assay quantified the protein concentration in the supernatant for the two lowest temperatures. The average of the two lowest temperatures was calculated and the volume equivalent to 30 µg of protein in the lowest temperature was moved from each temperature fraction into a clean tube. Following the distribution of protein, each tube was brought to a final volume of 100 µL by the addition of S-Trap lysis buffer with inhibitors. After acidifying the samples, they were loaded onto the S-Trap column and centrifuged at 4,000 x g for 1 min to trap the proteins. The column was washed with an S-Trap binding buffer to remove contaminants. Trypsin in 50 mM TEAB was added to the S-Trap column and incubated at 37 °C for 3 h. Peptides were eluted from the S-Trap column and dried using a vacuum concentrator. The dried material was reconstituted in 100 mM TEAB. TMT reagent was added to the peptide solution and incubated for 1 h at room temperature. The reaction was quenched with 5% hydroxylamine. Peptides were cleaned using a desalting column. The labeled peptides were fractionated using high PH reverse-phase chromatography to reduce sample complexity. The fractionated peptides were analyzed using LC-MS/MS, and MS2 was used for quantification and identification of peptides. Proteome Discoverer was used for quality control, statistical analysis, and differential expression analysis [66]. The differentially expressed proteins were also analyzed by Enrichr to obtain the significantly stabilized and destabilized pathways.

### Western Blot

The Western blot analysis was performed as described before [42]. After 48 h treatment of each compound, the cells were lysed with RIPA buffer (50 mM Tris-HCl pH 7.5, 0.1% SDS, 1% Triton X-100, 150 mM NaCl, 0.5% Sodium deoxycholate, and 2 mM EDTA) containing protease/phosphatase inhibitor cocktail (Thermo Fisher Scientific, Waltham, MA, USA) at 4°C for 1 h. The supernatants were collected and quantified using BCA protein assay kit (Thermo Fisher Scientific) according to the manufacturer’s instructions. To identify the molecular weight, we used a regular range protein marker and precision plus protein dual color standard marker. Protein of 10–12 µg was loaded on 4–12% sodium dodecyl sulfate polyacrylamide gel electrophoresis (SDS-PAGE), and then transferred to polyvinylidene difluoride membrane. The membranes were blocked with TBS-T buffer (Tris-buffered saline containing 0.1% Tween 20) containing 5% (w/v) skim milk powder for 1 h at 25°C, and then incubated with primary antibodies for 18 h at 4°C. After incubation with the secondary HRP-linked antibody for 1 h at 25°C, the membranes were detected using Clarity™ Western ECL Blotting Substrates (Bio-Rad, Hercules, CA, USA) and a Bio-Rad imager. Quantification of the images was performed using the Image J software.

### In situ quantitative imaging of cellular cholesterol

Spatiotemporally resolved *in situ* quantification of cholesterol in the inner leaflet of the plasma membrane (IPM) of mammalian cells was performed using a ratiometric cholesterol sensor, WCR-*e*Osh4 as described previously [67, 68]. WCR-*e*Osh4 was prepared and calibrated using giant unilamellar vesicles as described previously [67]. WCR-*e*Osh4 was microinjected into cells, and the cholesterol concentration in the IPM was determined using in-house programs written in MATLAB as described [67]. The three-dimensional display of the local cholesterol concentration profile was calculated using the Surf function in MATLAB.

### In vivo studies

Female ovary-intact athymic nude mice were purchased from Jackson Laboratory at 6 weeks of age. Animals were acclimated for a week upon arrival and then were subcutaneously (s.c.) inoculated in flanks with either MCF-7 cells (1 million cells/animal, 12 animals) or MDA-MB-231 (1 million cells/animal, 18 animals). After the xenograft tumors reached the palpable size of 0.8 cm in diameter, the daily administration (s.c) of LicA (80 mg/kg.day) or vehicle (4% DMSO, 6% EtOH + 45% water + 45% PEG-400) was started and continued until day 28 after the first dose. Animals were weighed once per week and the tumor size was measured twice per week using a digital caliper. The size of tumors was calculated using the formula [Tumor volume = (A^2^) x B/2] with “A” representing the small diameter and “B” representing the large diameter of the tumor. The tumor volume was plotted for each animal every time the tumor was measured. The growth of tumors in animals treated with LicA was compared with growth of tumors in the animals receiving the vehicle control. After the completion of the treatments, animals were sacrificed and the tumors were harvested, weighed, and preserved for future studies. It should be noted that if an animal met any humane endpoint criteria before the completion of the treatment course, as per the IACUC protocol, we sacrificed the animal. ***Ethics declaration:*** All the in vivo studies and procedures were performed under the approved Northwestern University IACUC protocol # <u>IS00013602</u>.

### Statistical analysis

The regular ordinary least square regression (eg. simple linear regression) does not consider heterogeneity across groups or time. Therefore, we fit a linear model using generalized least squares (GLS) to compare the tumor size trend over time between groups using LicA and vehicle for MDA-MB-231 and MCF-7 xenografts. We also compared the marginal tumor size between groups using LicA and vehicle for MDA-MB-231 and MCF-7 xenografts through least-squares means. We compared different variance-covariance structures, through ANOVA and selected the best fitted model with the smallest AIC.

## Results

### Licochalcone A alters gene expression and pathway activity in breast microstructures from high-risk postmenopausal women

We exposed microstructures produced from fresh surgical breast tissue of 6 postmenopausal women to LicA (5 µM) or DMSO for 24 h and performed total RNA-seq. LicA produces a transcriptional profile that is very different from that of DMSO. Differential gene expression analysis showed that in LicA-treated samples 2341 genes were downregulated (adj P < 0.05) with 381 genes having logFC < −1. Down-regulated genes include *HMGCR*, *MVD*, *MVK*, *SQLE*, *ACAT2*, *SREBF1*, *SREBF2*, *INSIG1*, *LSS*, *PTGST*, *RELA*, *ALDH1A3*, *SERPINB2*, *Wnt*, and *CYP1B1*. In these same samples, a total of 1462 genes were upregulated (adj P < 0.05) with 538 genes having logFC > 1. Top ranked genes associated with the upregulated pathways include *NQO1*, *NQO2*, *HMOX1*, *GST*s, *TXN*, *TXNRD1*, *SLC*s, *G6PD*, and *GCLC*. Pathway enrichment analysis was performed, and the top downregulated, and upregulated pathways are shown in Figure 1A and Figure 1B and in the associated supplementary data (S1A and S1B). The analyses shown in Figure 1A and Figure S1A revealed that metabolic pathways including lipids and cholesterol metabolism and biosynthesis, PI3K-AKT signaling, and pathways involved in inflammatory response are among the top downregulated processes. In addition, cholesterol and acetyl-CoA appeared as the predicted decreased metabolites (Figure 1A). In contrast, as demonstrated in Figure 1B and Figure S1B, NRF2-dependent antioxidant pathways, pentose phosphate pathway (PPP), glutathione metabolism, suppression of EGFR signaling, and ferroptosis were among the top upregulated pathways. Transcription factor analysis (Figure 1C) revealed that the profoundly upregulated transcription factor in LicA-treated microstructures is *NRF2* which is consistent with our previous findings [42] and the significantly downregulated transcription factors in these samples are proliferative factors *SP1* and *KLF4*, inflammation transcription factor *NF-kB* and its subunit *RELA*, and the lipogenic transcription factors sterol regulatory element binding factor (*SREBF*) *1* and *SREBF2*.

**Figure 1.**
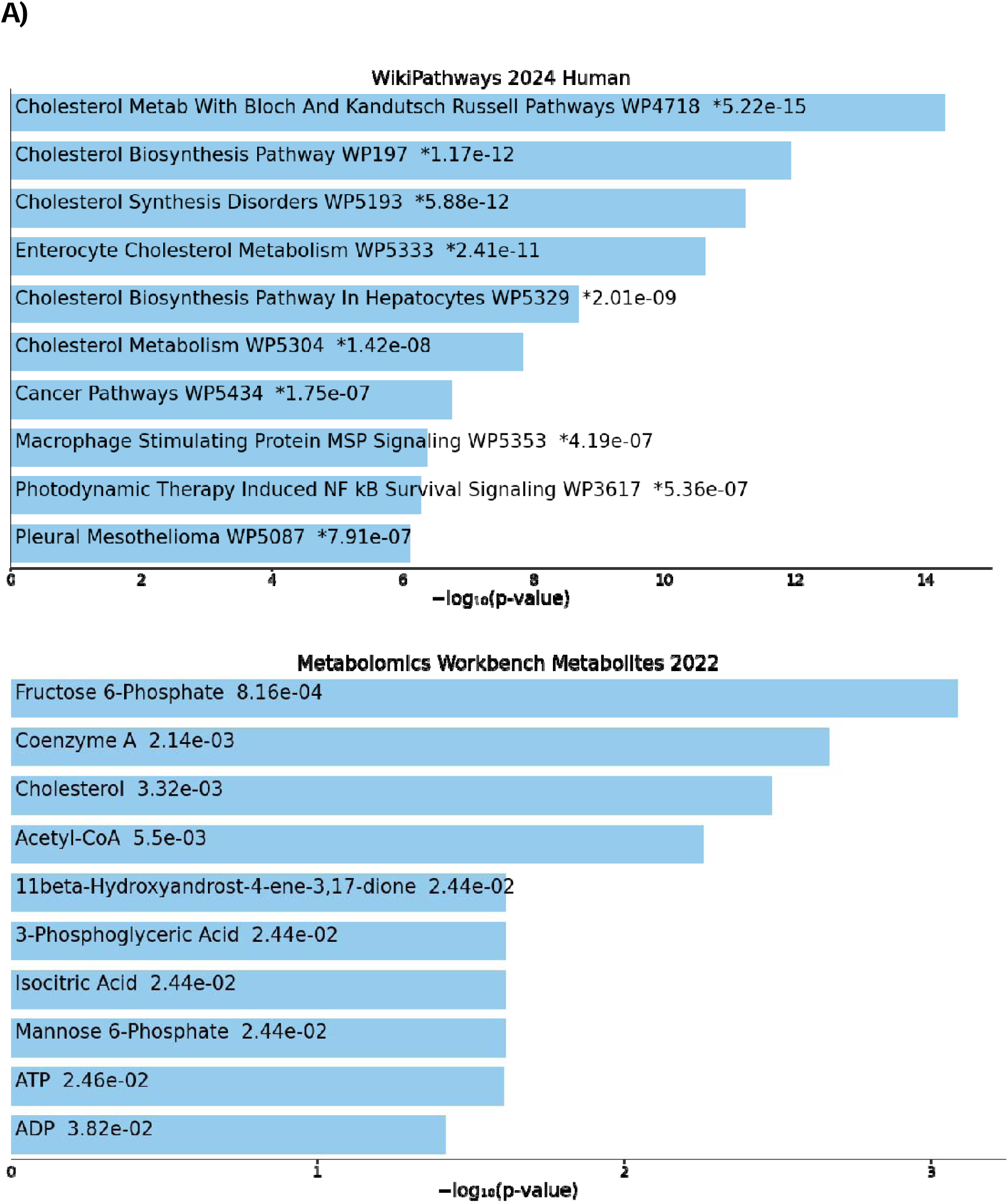
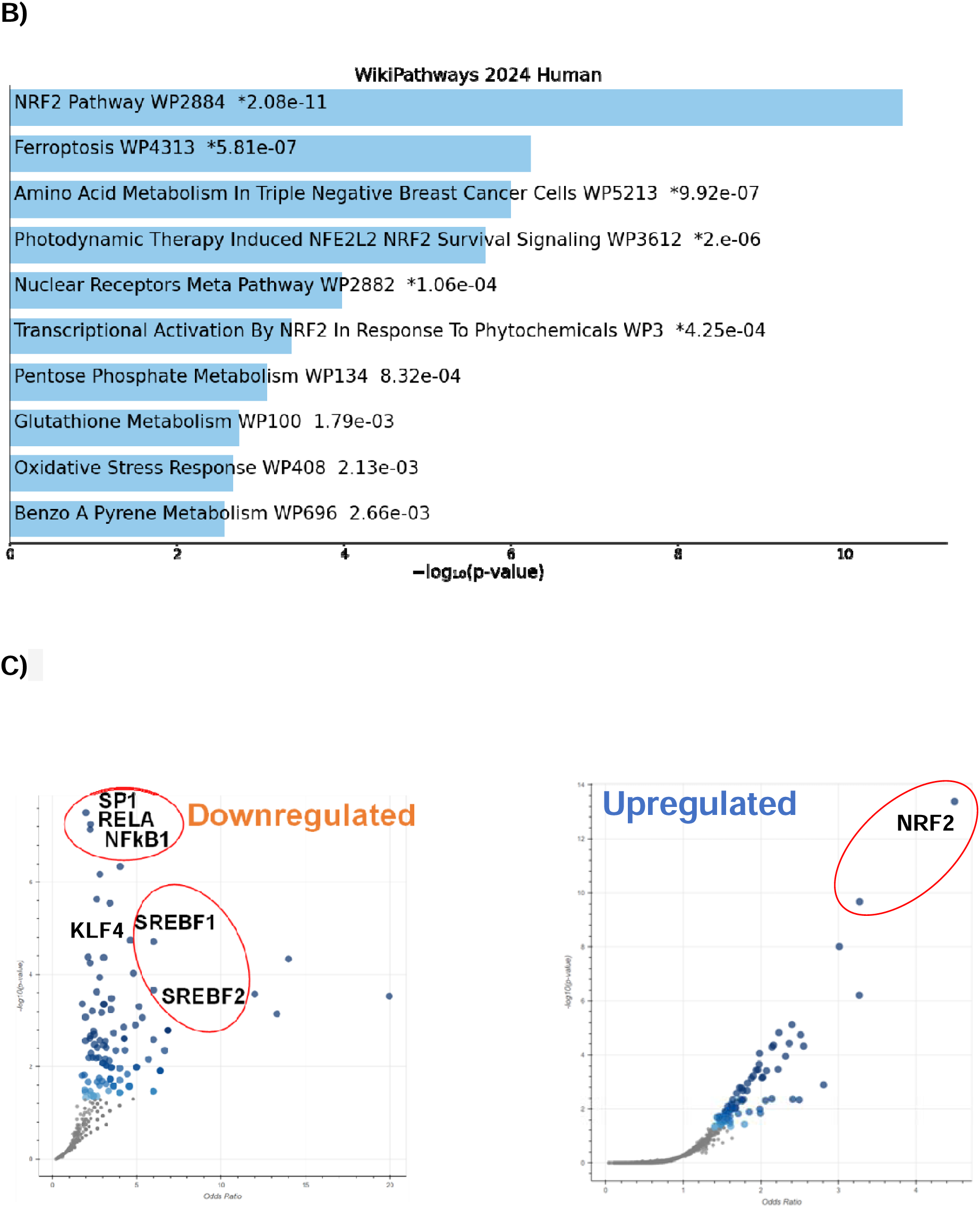
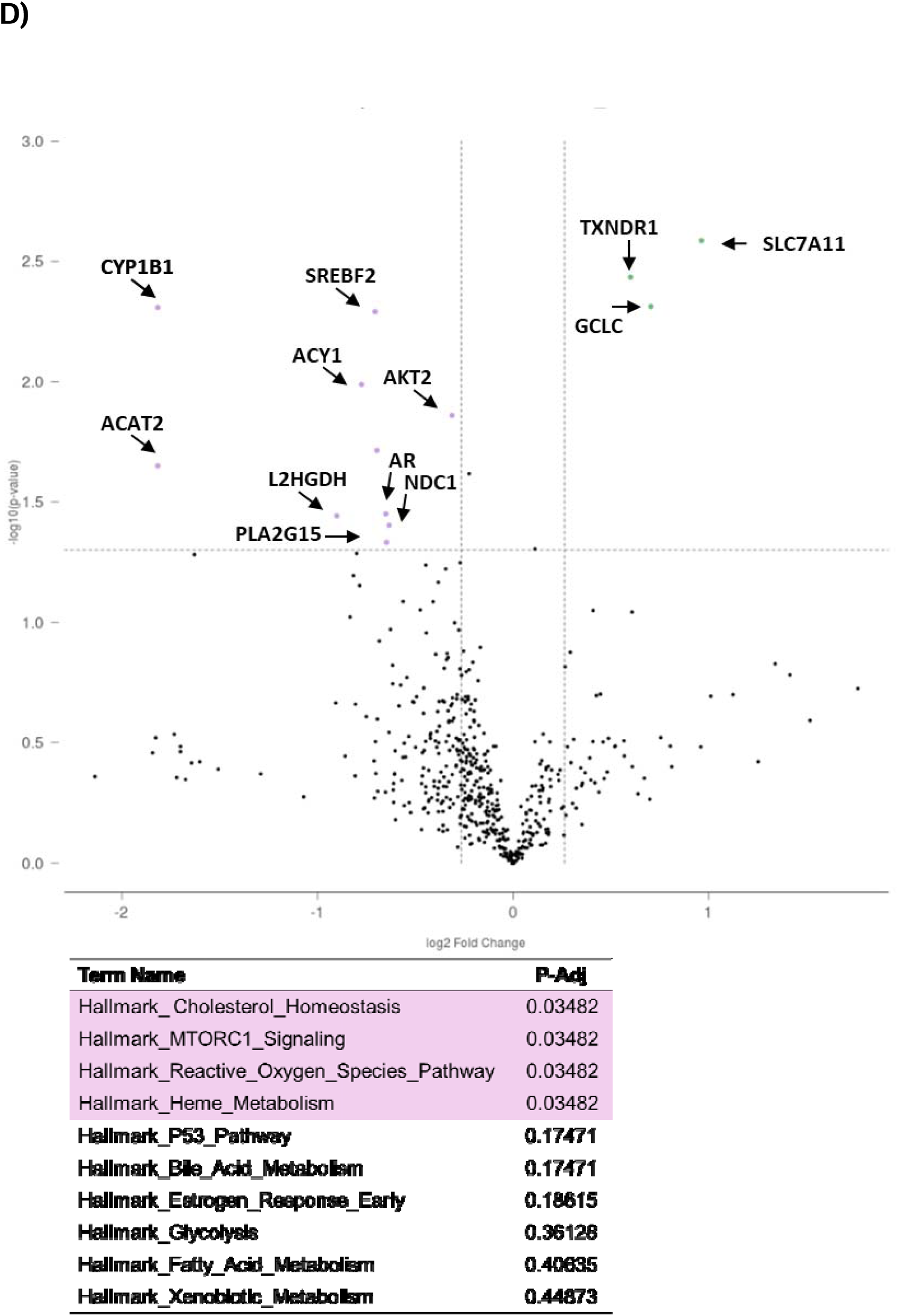
Transcriptomics analysis in postmenopausal high-risk women’s breast microstructures treated with LicA. Using RNA sequencing, LicA in the first set of 6 subjects’ specimens **A)** downregulates cholesterol and lipid metabolism, **B)** upregulates antioxidant pathways, **C)** downregulates transcription factors associated with inflammation, proliferation, lipid and cholesterol metabolism, and upregulates transcription factors associated with antioxidant response: data derived from TRRUST_Transcription_Factors_2019 gene set, **D)** shares significant similarities in downregulated and upregulated transcriptomic signatures with statins: data derived from drug matrix geneset. **E) NanoString metabolism panel analysis of six additional postmenopausal high-risk women’s breast microstructures**. LicA downregulates genes associated with lipid and cholesterol metabolism and estrogen carcinogenic metabolism and upregulates antioxidant response genes.

*<u>Validation of sequencing results</u>*, breast microstructures from 6 additional subjects were obtained. The NanoString metabolism panel consisting of 749 metabolic genes was employed and the results were analyzed using ROSALIND software. The data demonstrated significant (P < 0.05) downregulation of up to 3.5-fold change in *ACAT2*, *CYP1B1*, *L2HGDH*, *ACY1*, *SREBF2*, *NRF1*, *AR*, *PLA2G15*, *AKT2*, and *NDC1* expression and significant (P < 0.05) upregulation of up to 1.9-fold change in *SLC7A11*, *GCLC*, *TXNRD1* expression (Figure 1D). The analysis also showed that pathways associated with cholesterol homeostasis, mTORC1 signaling, reactive oxygen species metabolism, and heme metabolism were significantly modulated (adj P < 0.03) by LicA.

### Licochalcone A changes flux through metabolic pathways to facilitate support for antioxidant effects, reduced lipogenesis, and reduced proliferation

Further analysis of the sequencing data in the first 6 high-risk women’s tissue microstructures was performed to define flux through metabolic pathways using genome-scale metabolic modeling (Figure 2) [69, 70]. The analysis showed that flux through 191 reactions was significantly (p < 0.05) modulated as a function of exposure to LicA. Some of the top modulated metabolic reactions are key steps in glycolysis/gluconeogenesis, nucleotide biosynthesis and catabolism, the pentose phosphate shunt, pyruvate metabolism, cholesterol metabolism, fatty acid elongation, and steroid metabolism (Figure 2A). Interestingly, when the results were further explored at the metabolic reaction level using the BIGG database [71], the direction of the flux for many reactions was in favor of generating excess NAD(P)H and creating a reducing/antioxidant environment (Figure 2B). Examples of these reactions were aldehyde dehydrogenase in glycolysis/gluconeogenesis, glucose-6-phosphate dehydrogenase and phosphogluconate dehydrogenase in PPP, 17-beta hydroxysteroid dehydrogenase in steroid metabolism, and methylene tetrahydrofolate reductase in folate metabolism (Figure 2B). Flux through the NADPH-generating oxidative branch of PPP which is irreversible was significantly increased (Figure 2C). Flux through the non-oxidative branch of PPP which is reversible was toward the generation of ribose 5 phosphate (R5P) (Figure 2C). However, flux through PRPPS, which is responsible for *de novo* nucleotide biosynthesis, was not in favor of PRPP generation; rather the production of R5P was favored (Figure 2C) with the subsequent enhanced flux through PPM-catalyzed reaction which is involved in nucleotide catabolism and salvage production of nucleotides. Thus, the consumption of R5P and generation of R1P, which supports efficient recycling of nucleotides for DNA repair or energetics, is favored, as opposed to R5P formation essential for nucleotide *de novo* biosynthesis and cell proliferation (Figure 2C).

**Figure 2.**
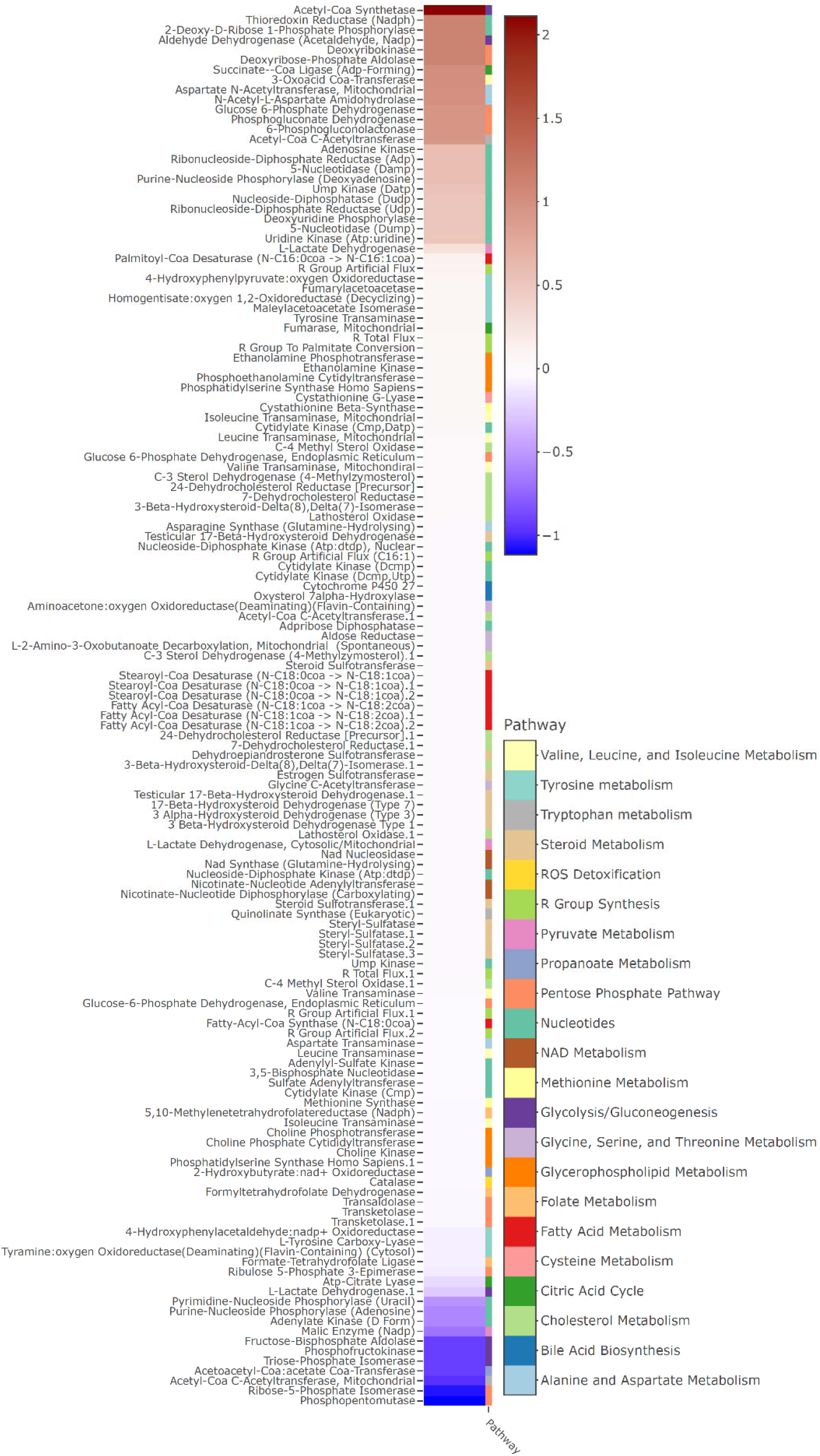
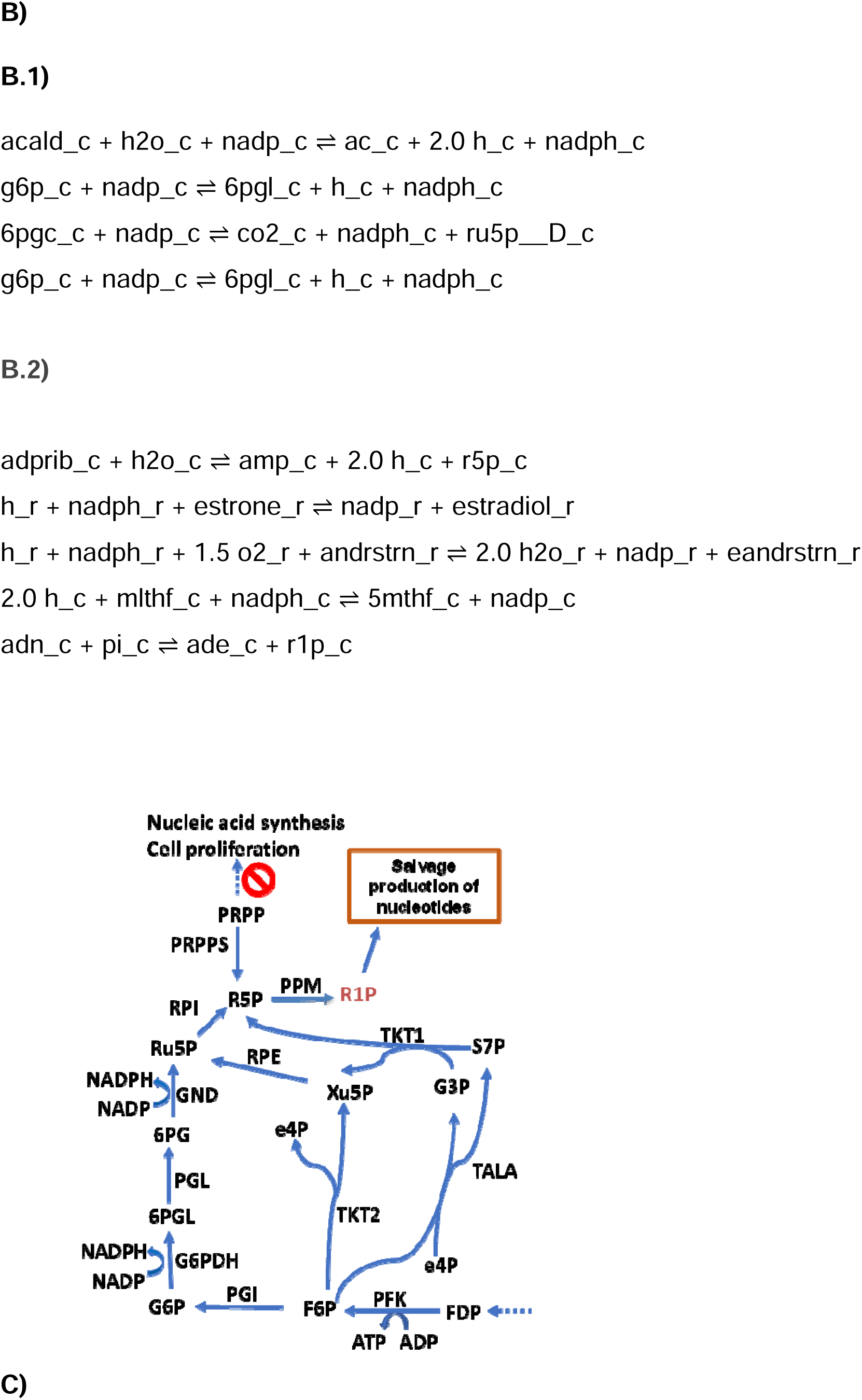
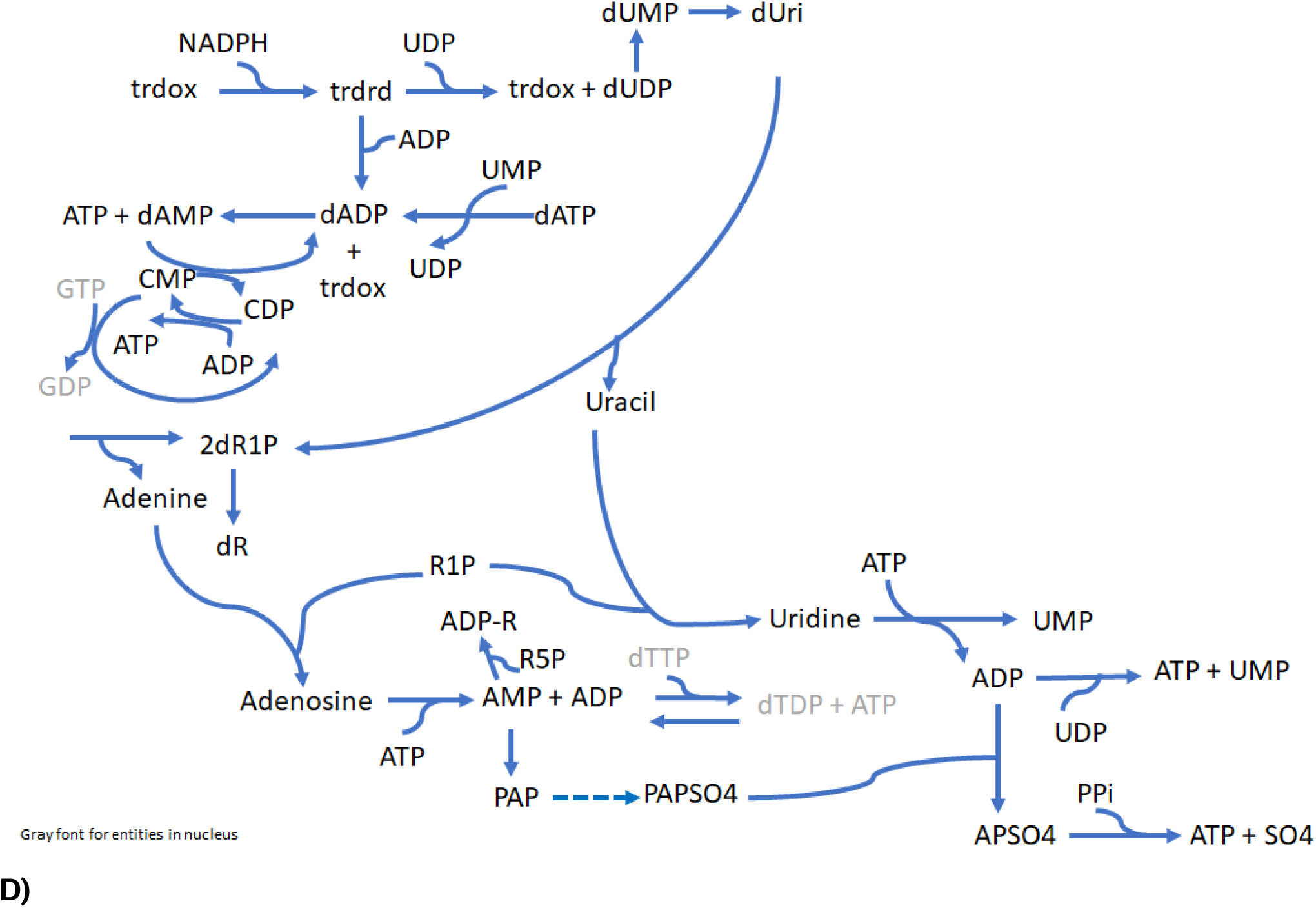
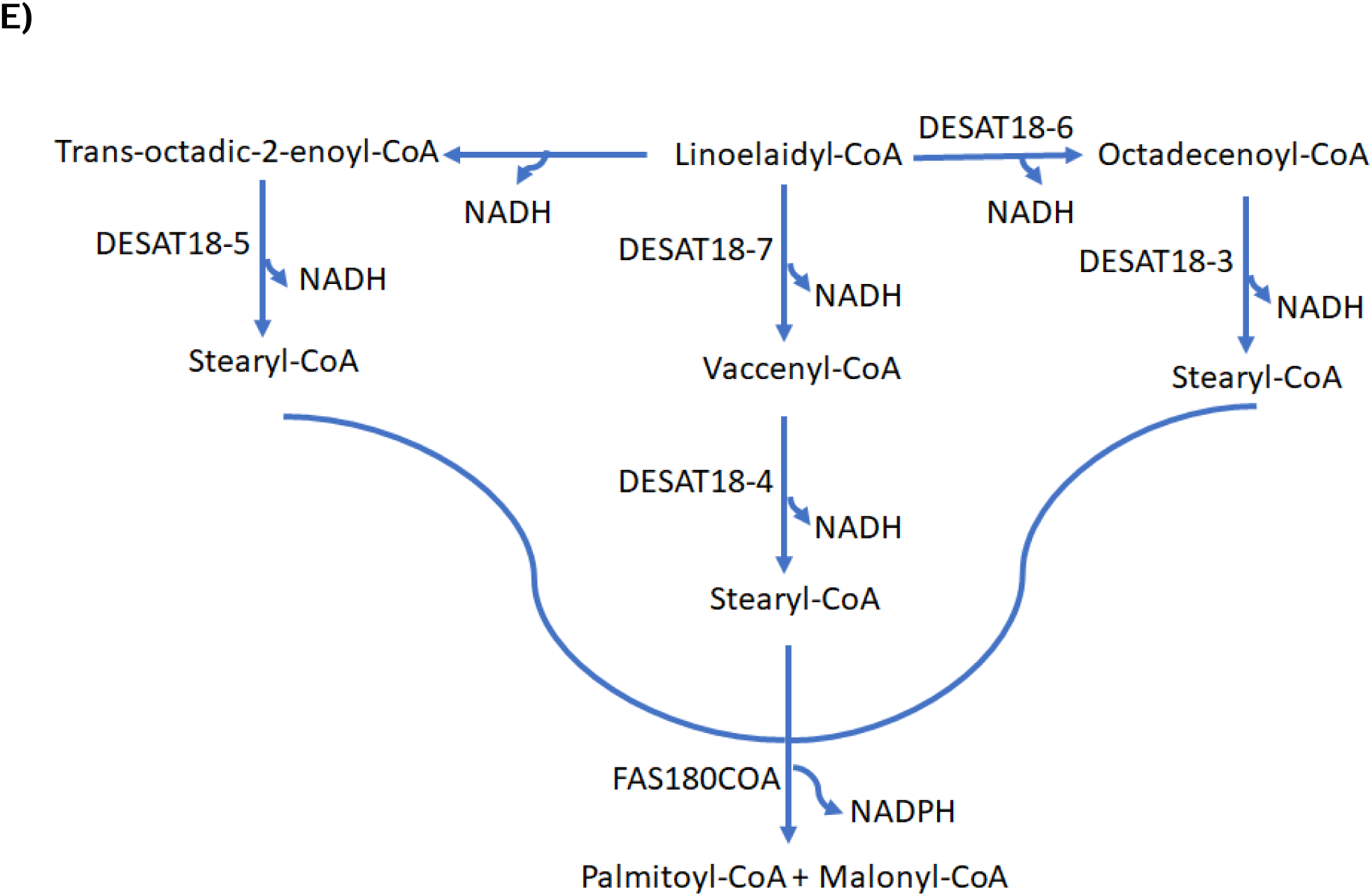
Genome-scale metabolic flux balance analysis in six postmenopausal high risk women’s breast microstructures. **A)** Metabolic flux heat map identifies significant flux (p-value < 0.01) through antioxidant pathways, lipid and cholesterol biosynthesis and metabolism. Magnitude of flux difference between treatment and control are shown. **B)** Metabolic flux in the representative pathways is (B.1) towards the right direction and (B.2) towards the left direction to enhance NAD(P)H generation. **C)** Metabolic flux through pentose phosphate shunt is in the direction generating NAD(P)H without enhancing nucleotide biosynthesis. **D)** Nucleotides essential for bioenergetics and kinetics of the metabolic reactions are salvaged. **E)** Flux through fatty acid elongation drives NAD(P)H generation and formation of saturated fatty acids.

*<u>Overall nucleotide flux</u>* (Figure 2D) offered further confirmation of the above findings. We observed the dominance of the salvage nucleotide biosynthesis process and its connection to antioxidant reactions such as NADPH-dependent formation of reduced thioredoxin (Figure 2D). This was further made evident when we observed that the formation of ADP-ribose from AMP and R5P is favored over the formation of R5P which can lead to reduced proliferation (Figure 2B and 2D). Similarly, we observed that flux through several reactions in cholesterol metabolism (Figure S2) such as 4-methyl zymosterol, 24-dehydrocholesterol reductase, 7-dehydrocholesterol reductase, lathosterol oxidase, cholesterol efflux, and several cholesterol transporters were in a direction to generate less cholesterol and excess NADPH (Figure S2). These were consistent with the differential gene expression results (Figure 1). In addition, flux through fatty acid elongation reactions catalyzed by desaturase enzymes such as stearyl CoA desaturase (SCD) were significantly modulated to enhance the ratio of saturated to monounsaturated fatty acids, and to generate excess NADPH, as a function of exposure to LicA (Figure 2E).

### Licochalcone A suppresses the proliferation of pre-malignant and malignant breast cells

Our transcriptomic data and metabolic flux modeling studies suggested LicA has antioxidant/anti-inflammatory effects along with decreased production of nucleotides which are required for cell proliferation. These observations encouraged us to evaluate how and at what concentration(s) LicA affects breast cell proliferation. We selected two pre-malignant breast cell lines; DCIS.COM (ER− PR-) and DCIS.COM (ER+ PR+), and 5 malignant cell lines MCF-7 (ER+ PR+), MCF-7aro (ER+ PR+ overexpressing CYP19A1), MDA-MB-231 (ER− PR-), HCC1937 (ER− PR-, BRCA1 mutation), and HCC3153 (ER− PR-, BRCA1 mutation). Exposing these cell lines to various concentrations (range of 350 nM – 40 µM) of LicA we observed that a single dose at 20 µM or 40 µM was able to exert a sustained suppression of proliferation in DCIS.COM and DCIS.COM/ER+ PR+ that lasted for at least 4 days post the single dose (Figure 3A). This observation held true for MCF-7 and MDA-MB-231 cells (Figure 3B, Figure 3C), which also responded to 10 µM of LicA, although the antiproliferative effect at this dose was not sustained following a single dose. Next, we tested repeated dosing of LicA, every 48 h for the maximum of 3 doses. Interestingly, two treatments of MDA-MB-231 cells with LicA (10 µM) 48 h apart until day 4 were able to sustainably retard proliferation until day 11 (Figure 3D) when we stopped our experiment. We also saw a reduction of proliferation of MCF-7 cells after 3 doses of LicA (10 µM) 48 h apart which was sustained at least until day 12, while MCF-7aro cells, HCC1937, and HCC3153 cells were also very responsive to LicA (5 µM) (Figures 3E - 3H). Our BrdU assay confirmed an anti-proliferative rather than a cytotoxic effect and the use of a ferroptosis inhibitor, ferrostatin, in combination with LicA showed that the antiproliferative effects are not due to ferroptosis (data not shown).

**Figure 3.**
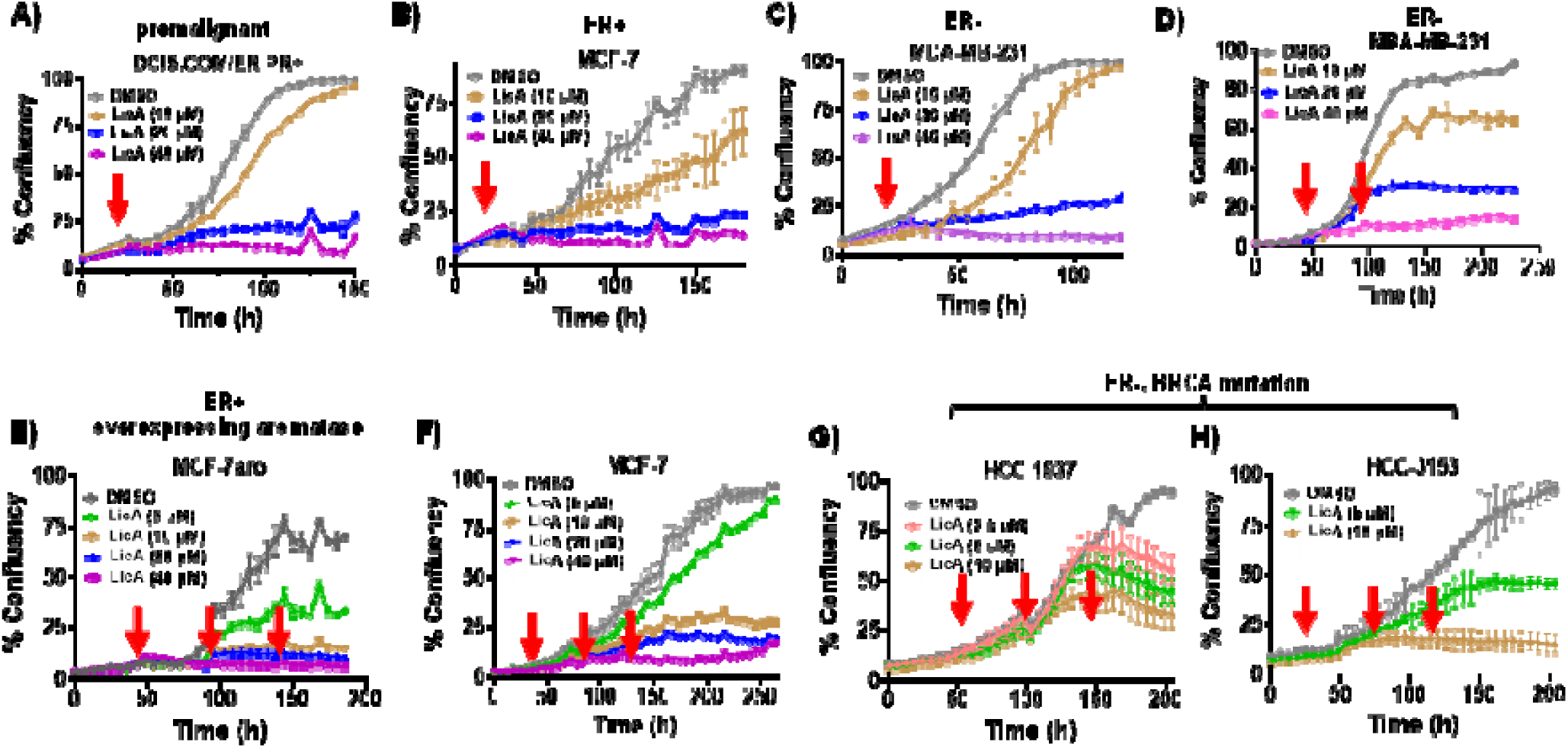
The effect of LicA on proliferation of ER+ and ER− premalignant and malignant breast cell lines. Repeated dosing of LicA exhibits sustained antiproliferative effects in aggressive cells. Red arrows show the time point(s) at which LicA or vehicle control was administered. Data represents mean ± SD of at least three independent replicates.

*<u>Supporting evidence for antiproliferative effects</u>* was provided with the NanoString data and the PISA proteomics results. Figure 4A demonstrates that the pro-proliferative genes such as *BCL2* in MCF-7 cells as well as *RRM2* and *MKI67* in *MCF-7* and MDA-MB-231 cells were significantly downregulated. Consistent with our metabolism flux data in breast microstructures which showed reduced *de novo* nucleotide biosynthesis, we also observed significant downregulation of *PRPS1* in MCF-7 cells (Figure 4A). Proteomics results (Figure 4B) further confirmed that in these cells, LicA destabilized enzymes essential for proliferation such as several EIFs, and the proliferation marker MKI67, and that the majority of destabilized proteins were associated with cell cycle and proliferation (Figure 4C).

**Figure 4.**
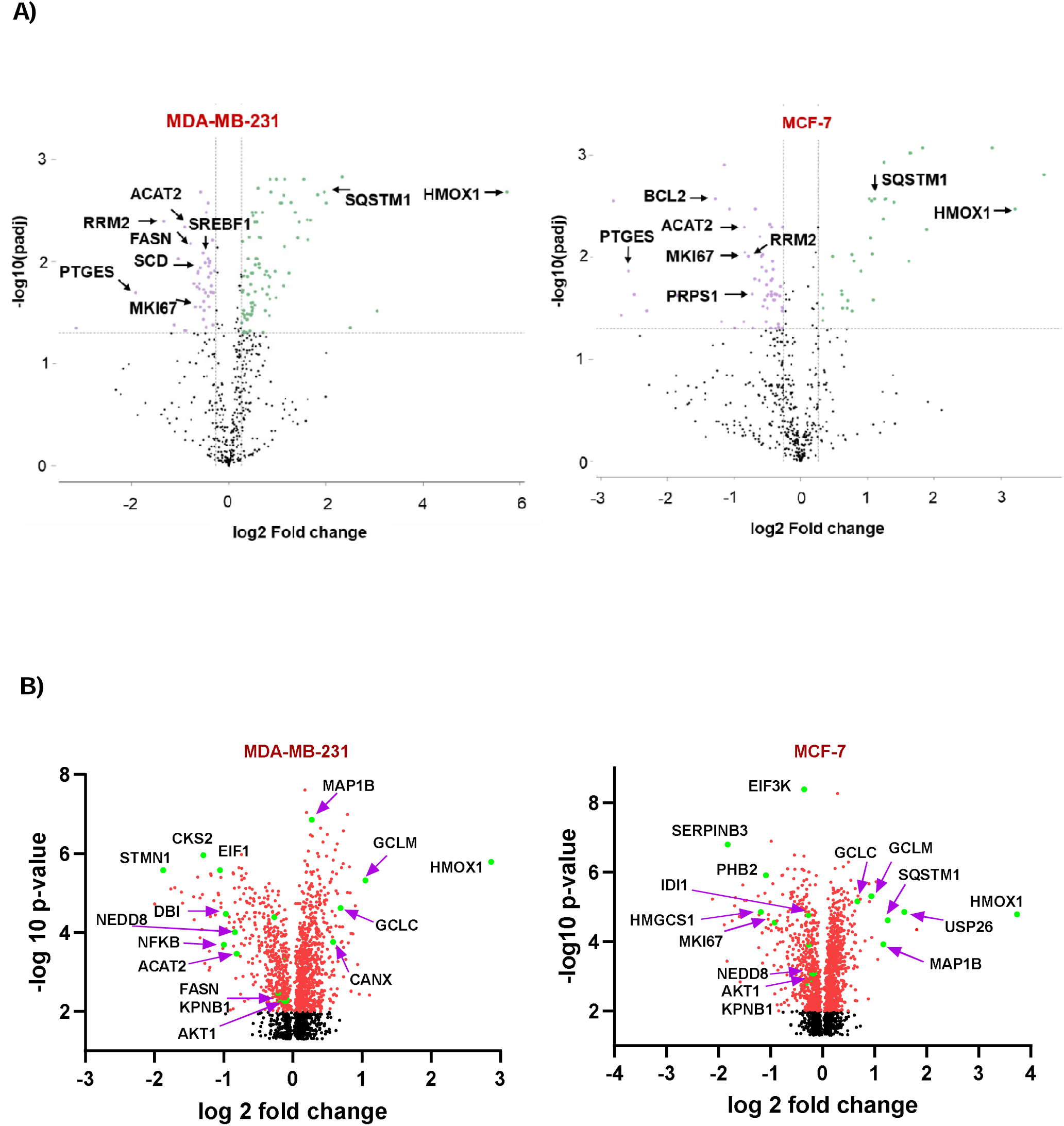
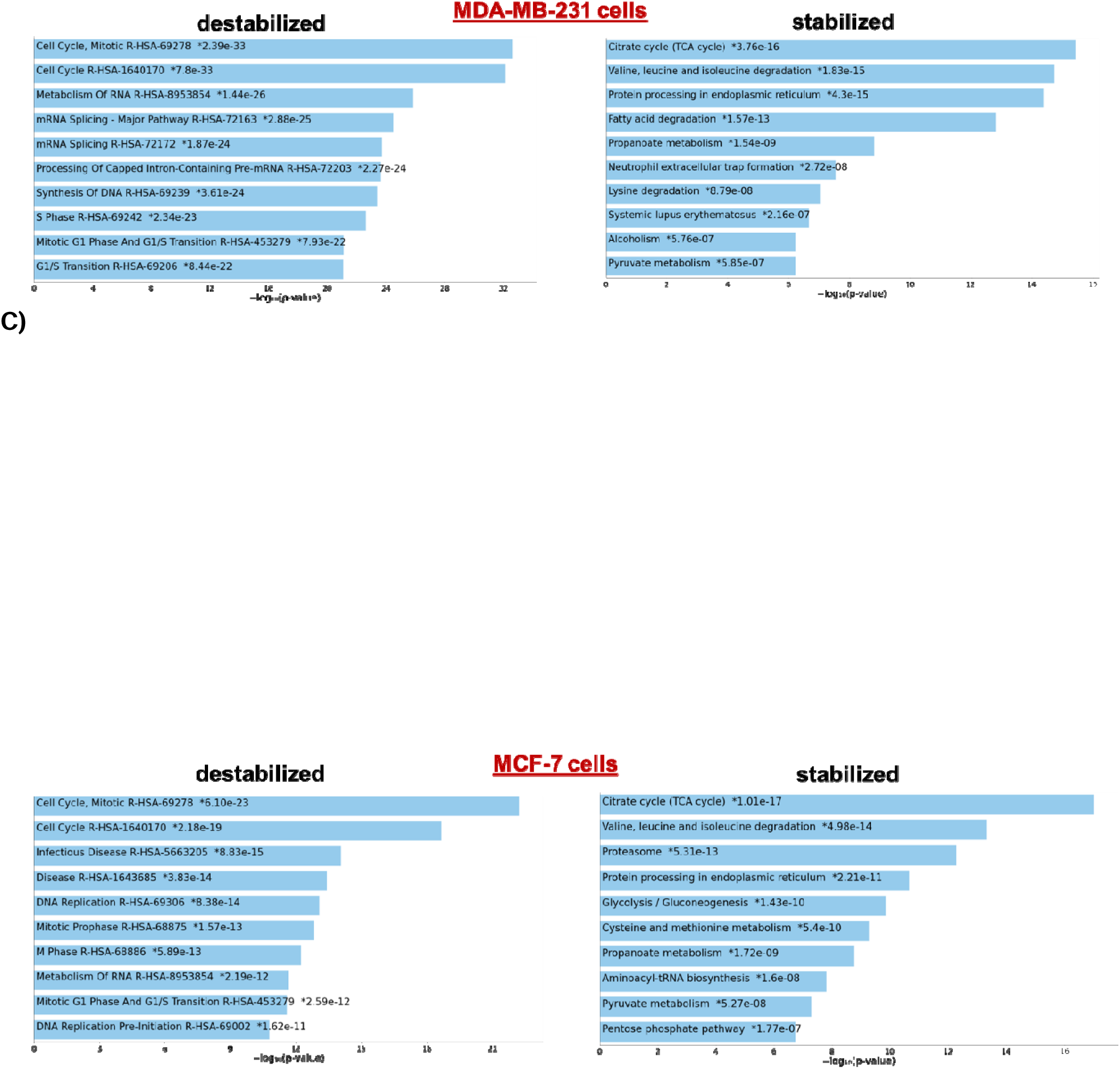
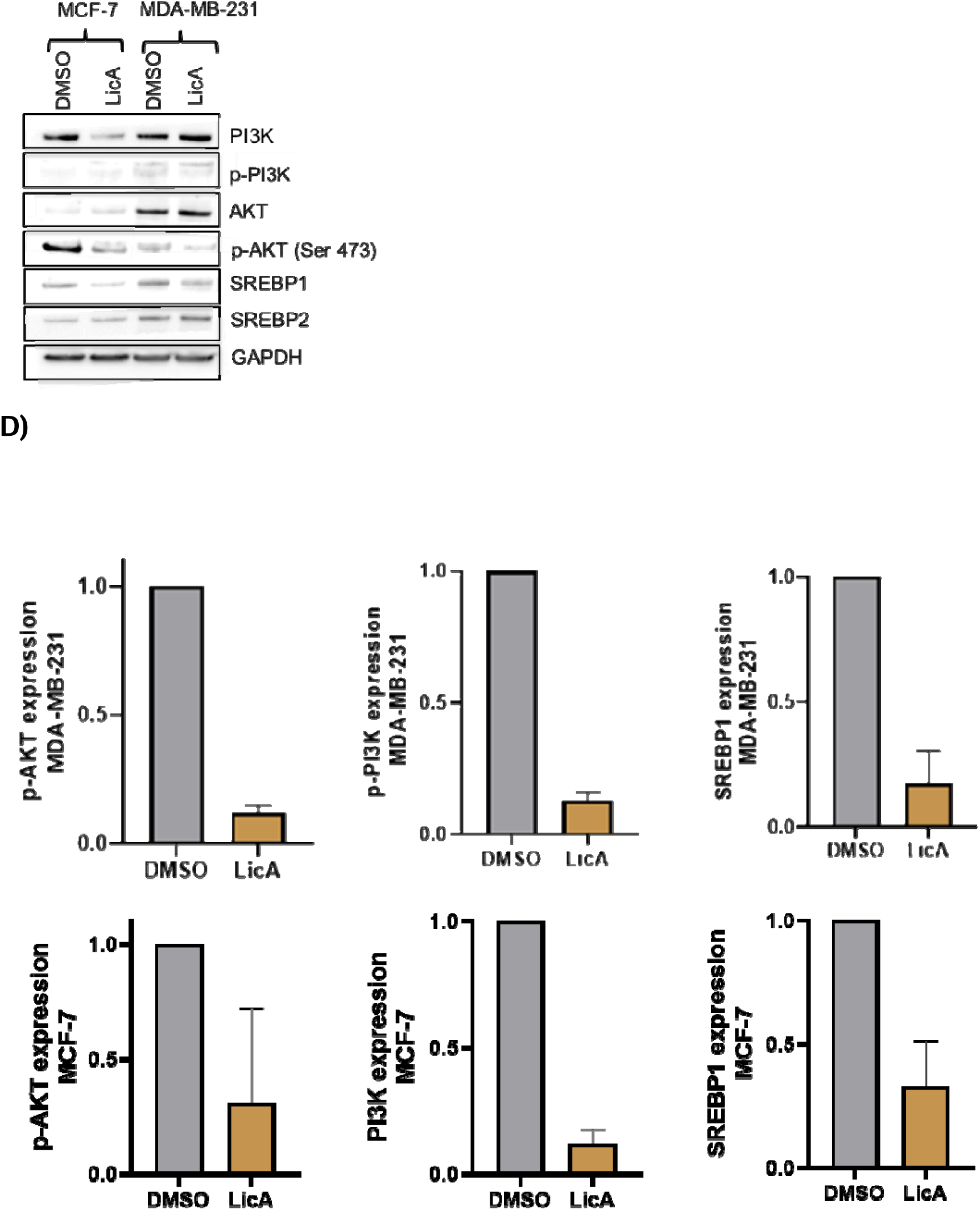
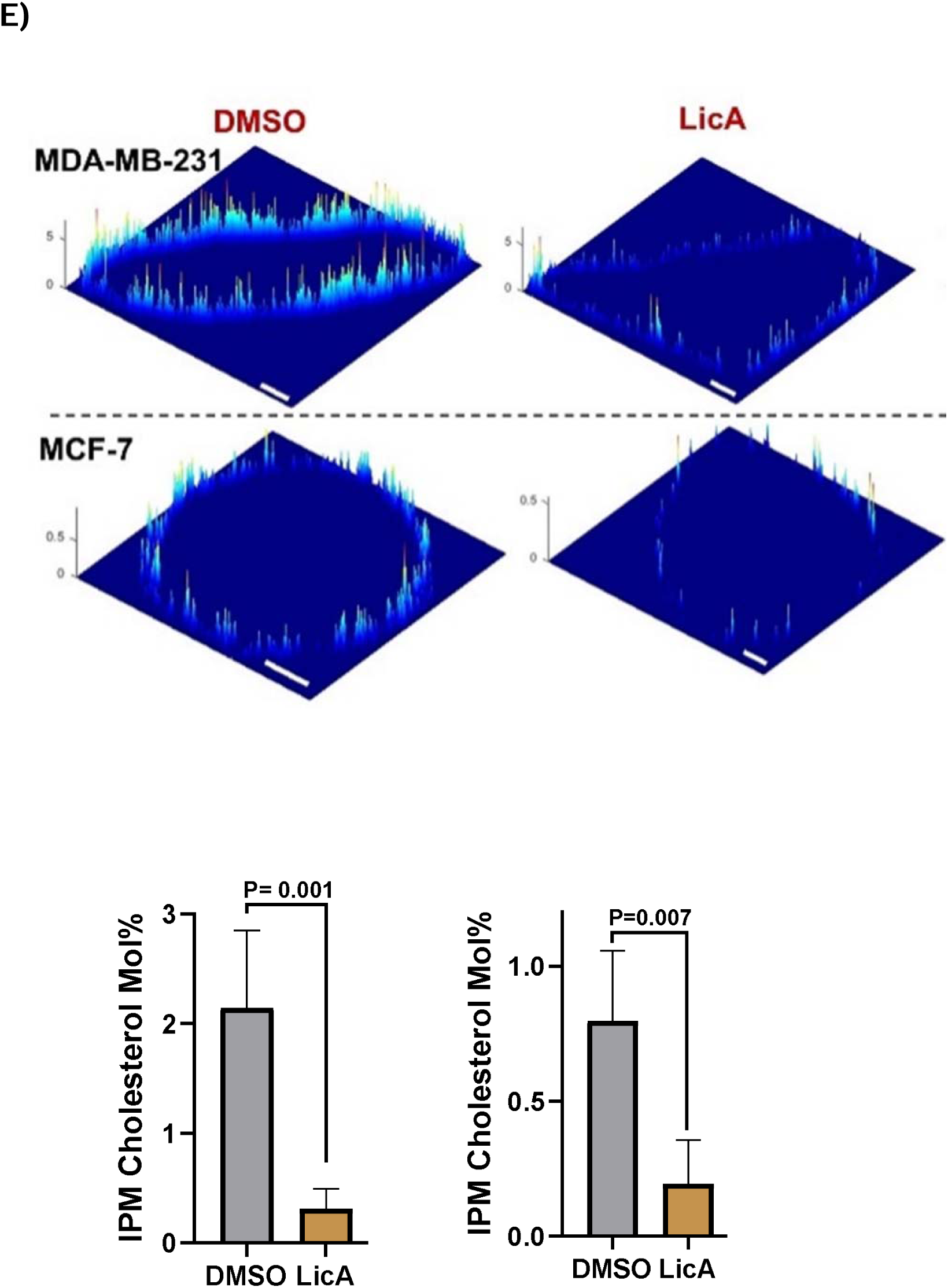
Changes in PI3K-AKT-SREBP1 lipogenesis in. MCF-7 (ER+) and MDA-MB-231 (ER-) treated with LicA (10 µM, 24 h). **A)** NanoString metabolism pathway panel analysis shows the downregulation of lipogenesis, pro-inflammatory, and pro-proliferative genes and upregulation of antioxidant and anti-inflammatory genes. Data represents the average of six analytical replicates of each cell line. **B)** PISA proteomics volcano plots demonstrate the proteins significantly stabilized or destabilized after the labeled extracted peptides were analyzed using LC-MS/MS. Data represents the average of six analytical replicates of each cell line and was filtered for the PSM numbers > 2 and the p < 0.05. **C)** Pathway identification using Enrichr based on the significantly stabilized and destabilized proteins observed in PISA output. **D)** Western blots: quantification of the bands was performed by Image J. Data represent mean ± SD of three independent replicates. **E)** Spatiotemporally resolved quantitative imaging of cholesterol in the inner leaflets of the plasma membranes (IPM). The IPM cholesterol concentrations before and after treatment with LicA (10 μM, 24h) were calculated from the two-channel cross-sectional ratiometric images of representative cells at steady-state. The *z*-axis scale indicates the cholesterol concentration (mol%). A pseudo-coloring scheme with red and blue representing the highest and the lowest concentration, respectively, is used to illustrate the spatial IPM cholesterol heterogeneity. Scale bars indicate 10 μm. A spatially averaged concentration (average ± SD from triplicate independent determinations with >10 cells per measurement) was calculated for each condition.

### LicA downregulates SREBP1-dependent lipogenesis and inflammation in MCF-7 and MDA-MB-231 cell lines

The antiproliferative effects in ER+ and ER− BC cell lines (Figure 3) with LicA, along with the metabolic effects detected in breast tissue microstructures (Figure 1) led us to test if the antiproliferative effects observed in cell lines could be related to changes in lipogenic metabolism and inflammation. Our NanoString metabolism panel results (Figure 4A) in MCF-7 and MDA-MB-231 cells were consistent with our observations in high-risk women’s breast microstructures (Figure 1) and clearly showed downregulation of SREBP1-dependent lipogenesis genes such as *ACAT2*, *FASN*, and *SCD* and the pro-inflammatory *PTGES* (PGE2 synthase), in agreement with the significant upregulation of anti-inflammatory *HMOX1*.

*<u>PISA proteomics results</u>* were consistent with our NanoString data and confirmed a clear separation of the LicA-treated and the DMSO-treated cells in both ER+ and ER− cell lines (Figure S3A and S3B). Destabilization of key proteins is shown in Figure 4B; these include lipogenic enzymes such as ACAT2, FASN, HMGCS1, and the inflammatory mediator NF-kB1 with a significant suppression of karyopherin β1 (KPNB1) which is needed for the translocation of NF-kB to the nucleus before its transcriptional activities. LicA also destabilized NEDD8 (Figure 4B) which is important for the stabilization of SREBP1 through the post-translational neddylation. On the other hand, the anti-inflammatory HMOX1 protein was profoundly stabilized (Figure 4B).

*<u>Analysis of proteomics results with Enrichr</u>* further demonstrated significant destabilization of cell cycle progression pathways (Figure 4C). Interestingly, while lipogenic metabolism was significantly destabilized, the TCA cycle, branched amino acids degradation, and fatty acids degradation appeared as the top stabilized pathways (Figure 4C) in both MDA-MB-231 and MCF-7 cells. Consistent with these observations, western blots (Figure 4D) demonstrated that SREBP1 expression, the phosphorylation of AKT at Ser 437, and the phosphorylation of PI3K were suppressed in both MDA-MB-231 and MCF-7 cells treated with LicA. While the levels of PI3K itself were reduced in MCF-7 cells, it had a slight but detectable increase in the MDA-MB-231 cells. The reduction in AKT phosphorylation was not accompanied by a change in AKT expression as a function of LicA treatment in these cell lines.

### LicA suppresses local cholesterol levels in MCF-7 and MDA-MB-231 cells

Transcriptomic data from the high-risk human breast microstructures and MCF-7 and MDA-MB-231 cells treated with LicA, along with our proteomics observations showed significant suppression of SREBP1, SREBP2, HMGCR with a shift in metabolism from lipogenesis to TCA cycle, suggesting that cholesterol biosynthesis could be affected by LicA. Cellular free (unesterified) cholesterol is mainly found in the plasma membrane in mammalian cells [72, 73]. We thus performed spatiotemporally resolved *in situ* quantification of IPM cholesterol in MDA-MB-231 and MCF-7 cells before and after LicA treatment [68]. The results showed that the spatially averaged IPM cholesterol levels were significantly different between the two BC cell lines (Figure 4E). Importantly, when these cells were treated with LicA, their IPM cholesterol levels were drastically reduced, i.e., 8-fold in MDA-MB-231 cells (n=14) and 4-fold in MCF-7 cells (n=19) (Figure 4E). These reduced IPM cholesterol levels were comparable to those reported for unstimulated primary mammalian cells [67], demonstrating the efficacy of LicA in IPM cholesterol depletion.

### Licochalcone A retards luminal and triple negative xenograft tumors in mice

Based on the antiproliferative effects of LicA in various breast cell lines (Figure 3) and the downregulation of the proliferation-related genes and pathways (Figure 4), we evaluated the effects of LicA in xenograft models of ER+ and ER− mammary cancers. In both xenograft models, analysis of results included all the animals assigned to each experimental group. In animals bearing MDA-MB-231 (ER-) xenografts, daily treatment with LicA (80 mg/kg.day, s.c.) for 28 days significantly (P < 0.000) reduced the growth rate of tumors in 7/9 (22%) of animals (Figure 5A). In the animals bearing MCF-7 xenograft tumors, treatment with LicA led to a significant (P < 0.005) reduction in the rate of tumor growth in all 6/6 (100%) animals, with a relatively similar responsiveness to treatment (Figure 5B).

**Figure 5.**
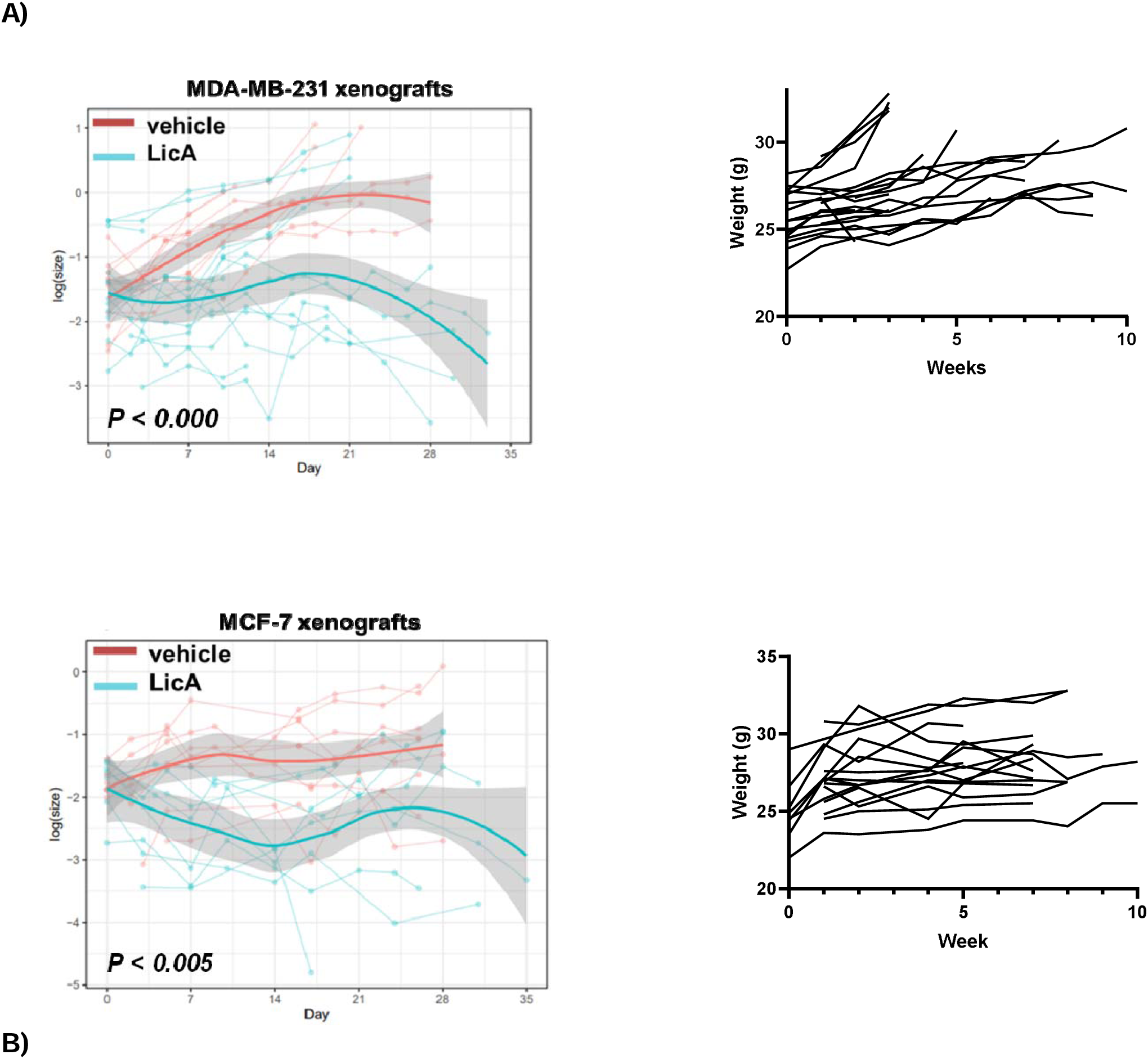
The effect of LicA on the rate of tumor growth in vivo. Xenografts of **A)** MDA-MB-231 (n =18 final tumor bearing animals) or **B)** MCF-7 cells (n = 12 final tumor bearing animals) were performed in the right (Rt) and left (Lt) flanks of ovary intact athymic nude mice. After the tumors reached 0.8 cm in diameter, subcutaneous administration of LicA (80 mg/kg.day) commenced and continued for 28 days. A significant reduction in the rate of tumor growth was observed in the LicA treated versus vehicle treated animals.

## Discussion

One of the hallmarks of cancer and an essential step in malignant transformation is metabolic reprogramming and changes in cellular bioenergetics that facilitate the high energy demand of hyperproliferating cells and promote their survival [39]. A main mechanism for this effect is the reprogramming of lipid metabolic pathways [74–79]. Within this major pathway, the *SREBF1* gene encodes SREBP1a and SREBP1c proteins which are transcription factors that regulate lipogenesis. *SREBF1* expression is significantly higher in breast tumors compared to normal tissue adjacent to tumor and in healthy breast tissue of 924 patients in the GTEx and TCGA datasets [80–83]. Conversely, the downregulation of SREBP1 not only decreases lipogenesis, but is also associated with bone protection [84–86]. It is known that SREBP1 regulates the expression of lipogenic factors, fatty acid synthase (FASN), acetyl-CoA acetyltransferase 2 (ACAT2), acetyl CoA carboxylase 1 (ACC1), and ATP citrate lyase (ACLY) which are essential for the survival of BC cells [25, 87, 88]. There is also a strong association between breast tissue stiffness (a risk factor for breast carcinogenesis) and SREBP1 activity [26, 89, 90]. SREBP1 activates NF-kB-dependent inflammation by promoting fatty acid desaturation [91, 92]. Recent studies have demonstrated that activation of NF-kB-dependent inflammation happens in part through the translocation of the inactive SREBP1-NF-kB complex to the Golgi apparatus and the subsequent activation of SREBP1 mediated by the SREBP cleavage activating protein (Scap) [28] suggesting a close connection between SREBP1 metabolism and NF-kB inflammation. An inflamed environment supports the growth of malignant cells through the recruitment of macrophages, secretion of inflammatory cytokines, enhancement of fatty acid oxidation, formation of reactive oxygen species (ROS), change of the hormonal milieu, and increased proliferation [93]. Upstream of SREBP1, EGFR activation and PI3K-AKT-mTOR mutation, as well as mTORC1 activity lead to enhanced SREBP1 expression and consequent increase in SCD activity known to be important for cancer initiation, promotion, and invasion [94–101].

We used breast microstructures from the opposite breasts of women with breast cancer, since these represent normal/benign tissue that is known to be at increased risk of a second new breast cancer. LicA significantly lowered *SREBF1*, as well as the expression of several of its upstream effectors such as *EGFR*, *PI3K*, *AKT*, and *mTORC1*, in addition to *SCD* which is downstream of *SREBF1*. These observations extended to MDA-MB-231 and MCF-7 cells, where transcriptomics and proteomics data clearly demonstrate significant reduction of lipogenesis signals regulated by SREBP1, in concert with a significant anti-inflammatory response with suppression of PGE2 biosynthesis and a profound increase in HMOX1 expression. Interestingly, SREBP1 has response elements at the promoter region of *HMOX1*, i.e. downregulation of SREBP1 is expected to lower *HMOX1* expression; however, in our results, HMOX1 at gene and protein levels is profoundly enhanced. This could be associated with the effect of LicA on the KEAP1-NRF2 pathway [42] which activates HMOX1 and compensates for its likely reduction due to SREBP1 downregulation (Figure 6). In our results, the significant decrease in the expression of SREBP1 is accompanied by reduced expression of NEDD8 protein which is important for the stabilization of SREBP1 through posttranslational neddylation, a process linked to BC aggressiveness [102]. In addition, lowered SREBP1 expression along with the reduced phosphorylation of its upstream effectors PI3K and AKT and the profound drop in cellular cholesterol levels is aligned with the metabolic shift observed in our proteomics studies. The combined effects of LicA on stabilizing TCA cycle, degradation of branched chain amino acids, and fatty acids along with significant destabilization of the cell cycle show that LicA reprograms cellular metabolism in favor of reducing cell proliferation [103, 104]. These results demonstrate that the axis of PI3K-AKT-SREBP1-NF-kB is the mechanism by which LicA exerts its antiproliferative and tumor-suppressive effects. LicA’s precise target in this axis is yet to be elucidated.

**Figure 6.**
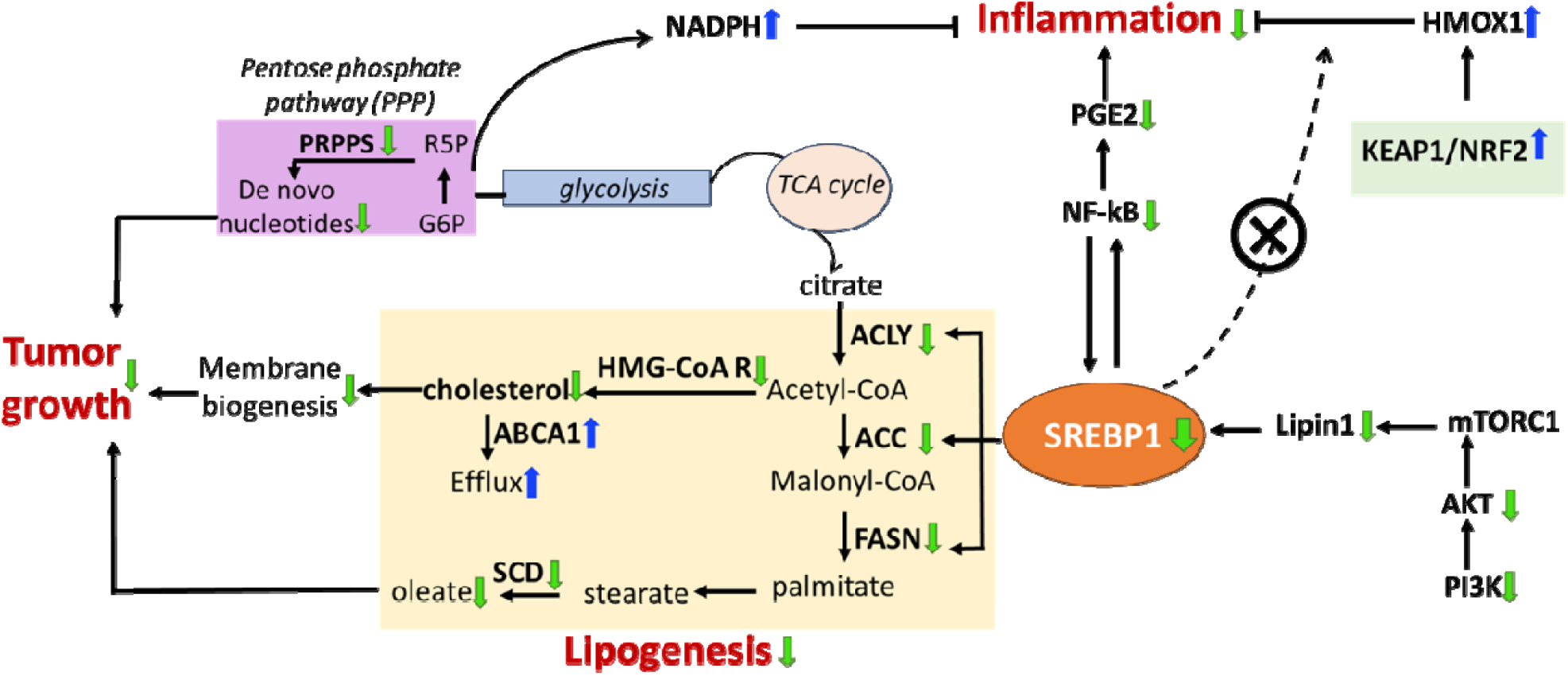
Suppressing lipogenesis and inflammation, mediated by the central role of SREBP1 regulation by LicA, leads to reduced proliferation in the breast. Colored arrows represent the effects of LicA.

Another important gene in lipid metabolism is the *SREBF2* encoding the SREBP2 protein which is under the control of SREBP1 and primarily regulates cholesterol biosynthesis and homeostasis [105]. It is known that enhanced expression of *SREBF2* promotes the progression of several types of cancers [76, 106, 107]. *SREBF2* is significantly higher in breast tumors compared to healthy tissues of 924 patients in GTEx and TCGA samples [80–83]. Among the genes that are regulated by SREBP2 are members of the mevalonate pathway such as *MVD*, *MVK*, *SQLN*, and *HMGCR*. The activity of *SREBF2* is opposed by the major tumor suppressor P53 through transcriptional upregulation of the ABCA1 transporter and cholesterol efflux [105]. This effect is further reinforced by the activation of LXR and its subsequent regulation of cholesterol homeostasis and ABCA1 mediated efflux, suppressing *de novo* biosynthesis, and uptake of cholesterol. In our study in women’s breast tissue in addition to lowered *SREBF2* expression, the effect of LicA on the Mevalonate Pathway resembles the effects of p53 and LXR activation. This once again shows that LicA reverses the oncogenic direction of the EGFR-PI3K-AKT-SREBP axis in non-malignant tissue, and thus can protect breast cells by reprogramming metabolism. It was recently reported that cholesterol ([Chol]_I_) in the inner leaflet of the plasma membrane (IPM) of mammalian cells is directly and quantitatively related to the proliferative activity of cells [68]. The significant reduction in cholesterol biosynthesis in the inner leaflet of plasma membrane which we depicted by measuring [Chol]_I_ in MCF-7 and MDA-MB-231 cells further validated the metabolic reprogramming. The difference we found in the [Chol]_I_ between MCF-7 and MDA-MB-231 cells reflects differential reprogramming of cholesterol homeostasis in these two cell lines.

Interestingly, changes in the SREBP2 expression in women’s breast microstructures and in malignant breast cell lines MCF-7 and MDA-MB-231 differ. In women’s breast microstructures which are nonmalignant and composed of a mixture of various cell types in breast tissue SREBP2 was profoundly downregulated when the tissue was exposed to LicA; however, in malignant breast cell lines there was not a significant change in SREBP2 levels at the RNA and protein levels, although IPM cholesterol was significantly suppressed. Thus, LicA reduces lipogenesis in non-malignant high-risk breast through reducing the expression of both SREBP1 and SREBP2 while its effects on BC cells are primarily mediated by SREBP1. A recent study showed that concerted SREBP1-NF-kB modulation might exert metabolic and inflammatory responses independent of SREBP2 levels or activity, as it is executed through the recruitment of SREBP1 and NF-kB to the Golgi apparatus mediated by and in complex with Scap [28]. In other words, it is highly likely that the reduced proliferation in cancer cells are primarily regulated by the SREBP1-NF-kB complex, through direct interaction with Scap. However, in non-malignant high-risk breast tissue the additional reduction in SREBP2 levels, reinforces prevention of cellular transformation. It is also likely that despite the tumor suppressive effects of LicA in both MCF-7 and MDA-MB-231 xenografts, the precise LicA target(s) could be cell type-dependent, since there were differences in SREBP1, PI3K, and AKT protein level changes in these cell lines.

We found that metabolic flux in high-risk women’s breast exposed to LicA was not only in favor of generating excess NADPH to support antioxidant and anti-inflammatory pathways, flux was also reprogrammed to minimize cell proliferation; the effect that we were able to confirm in a series of experiments including proliferation assessments, *in vivo* tumor growth studies, and molecular markers at gene and protein levels.

In addition to the SREBP1 dependent reduction in lipogenesis and inflammation that lead to reduced cell proliferation, our data once again shows LicA limits oxidative and inflammatory stress through the activity of NRF2 and suppressing NF-kB pathways in the breast tissue microstructures of medium-risk women. Such stressors can cause DNA damage, remodel extracellular matrix, promote cancer cell stemness, and modify paracrine effects to initiate and promote tumors [55, 93, 108–111]. Moreover, the significant expression of SQSTM1 in the breast microstructures, as well as in MCF-7 and MDA-MB-231 cells treated with LicA supports its previously reported bone protective effects [52–54].

LicA-based interventions will effectively reduce BC risk (Figure 6). LicA protects against oxidative and inflammatory stress, deprives cancer stem cells of their favored inflammatory environment, while it lowers lipogenesis and proliferation. LicA has compliance with Lipinski’s rule of 5 (RO5) in drug discovery; it has 2 hydrogen bond donors (RO5 standard < 5), has 6 hydrogen bond acceptors (RO5 standard <10), exact molecular mass = 338.15180918 g/mol (RO5 standard < 500 g/mol), and a calculated octanol-water partition coefficient = 4.9 (RO5 standard < 5). It should be noted that Lipinski’s RO5 requires a drug-like compound to comply with at least 3 out of 4 rules to remain orally active. Based on these physical properties and the biological activities reported herein and previously, LicA is a good candidate for further development as a novel BC risk reduction drug with sufficient efficacy, reduced toxicity, and therefore anticipated to have greater acceptance by women at increased risk. Such an agent with effects against both ER+ and ER− subtypes, and demonstrated bone protective properties is predicted to have a better success and a larger impact than the current agents. We are now planning the next series of studies including evaluations in immunocompetent models of cancer prevention, and rigorous formulation and pharmacokinetic evaluations for oral delivery, that are required for translation to clinical testing.

## Acknowledgments

We thank the Department of Surgery at Northwestern University and the Robert H Lurie Comprehensive Cancer Center for their support of this study. We also thank Dr. Xiaoling Xuei from the Indiana University Center for Medical Genomics for her advice during the RNA sequencing studies, Dr. Marcus Peter from Northwestern University for giving us access to IncuCyte, Dr. Shiuan Chen from the Beckman Cancer Center of the City of Hope for gifting us their MCF-7aro cells, and Dr. Dean Edwards from Baylor College of Medicine for gifting us their DCIS.COM/ER+ PR+ cell line.

## Author’s contributions

AH developed the hypothesis, designed the study, executed the in vitro, ex vivo, and in vivo studies, analyzed the data, wrote the manuscript. ETB conducted bioinformatics analyses. CHC and SC conducted the metabolism flux analyses and edited the manuscript, XG, KB, WC conducted the spatiotemporal cholesterol measurements, analysis, and edited the manuscript, OL assisted with the in vivo studies, RC analyzed in vivo data and provided biostatistical support. SEC supervised the study design and interpretation of data, edited the manuscript. SAK supervised the study design, clinical relevance, and interpretation of data, and edited the manuscript.

## Ethics approval and consent to participate

Institutional Review Board (IRB) approval (IRB STU00202331) was obtained from Northwestern University prior to obtaining informed consents from patients and collecting samples. All experiments were conducted in accordance with the approved protocol and guidelines. All the in vivo studies and procedures were performed under the approved Northwestern University IACUC protocol # IS00013602.

## Consent for Publication

Not Applicable

## Data availability

The datasets generated and/or analyzed during the current study are available in GEO repository, [PERSISTENT WEB LINK TO DATASETS]

## Competing interests

The authors declare that they have nothing to disclose and no competing interests.

## Funding

American Cancer Society postdoctoral fellowship 131667-PF-18-049-01NEC to AH, Robert H Lurie Comprehensive Cancer Center Translational Bridge Fellowship to AH, National Cancer Institute T32 postdoctoral fellowship to AH, Northwestern University Department of Surgery Seed funding to AH, Bramsen-Hamill Foundation funding to SAK, The National Institutes of Health grant R35GM122530 to WC.

## Notes

### Competing Interest Statement

The authors have declared no competing interest.

### Summary of Updates

Figure 1 revised Figure 4 revised Figure 5 revised Figure 6 revised Figures were reorganized to give a total of 6 figures The manuscript abstract and body were revised to reflect the additional information provided in the revised data figures. Additional authors were added.

