## Supplementary Data for "Reprogramming SREBP1-dependent lipogenesis and inflammation in high-risk breast with licochalcone A: a novel path to cancer prevention"

^1^Department of Surgery, Feinberg School of Medicine, Northwestern University, Chicago, IL. USA, ^2^Robert H. Lurie Comprehensive Cancer Center, Northwestern University, Chicago, IL. USA, ^3^Department of Biochemistry and Molecular Genetics, Northwestern University, Chicago, IL. USA, ^4^Department of Preventive Medicine, Northwestern University, Chicago, IL. USA, ^5^Department of Biomedical Engineering, University of Michigan, Ann Arbor, MI. USA, ^6^Department of Chemistry, University of Illinois Chicago, IL. USA.

***Corresponding Author:** Atieh Hajirahimkhan, PhD

303 E. Superior, 4-220, Chicago, IL. 60611

ORCiD identifier: 0000-0003-0585-5668

**S1A) Pathways upregulated in microstructures obtained from high-risk women’s breast tissue and exposed to LicA ex vivo.** Microstructures from 6 subjects were treated with LicA (5 µM) for 24 h followed by RNA sequencing, differential gene expression, and pathway analysis.

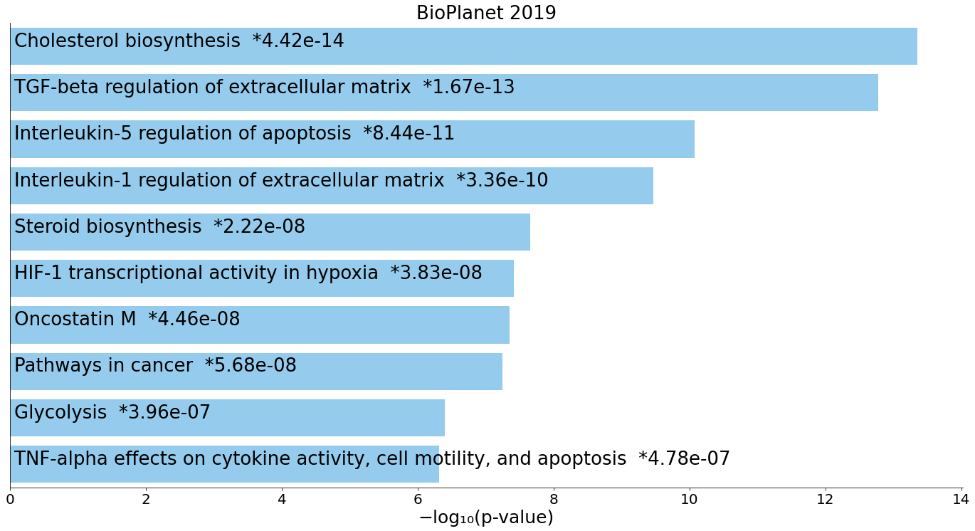

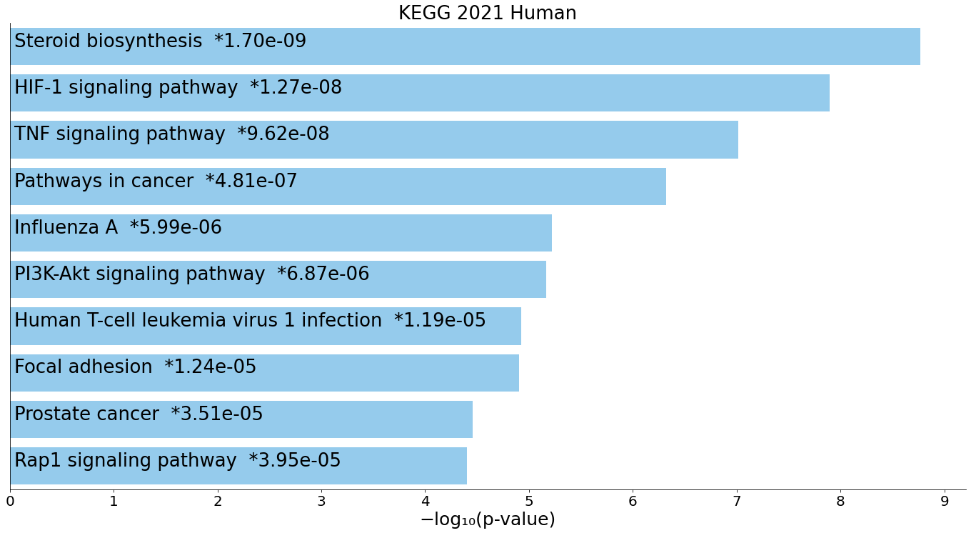

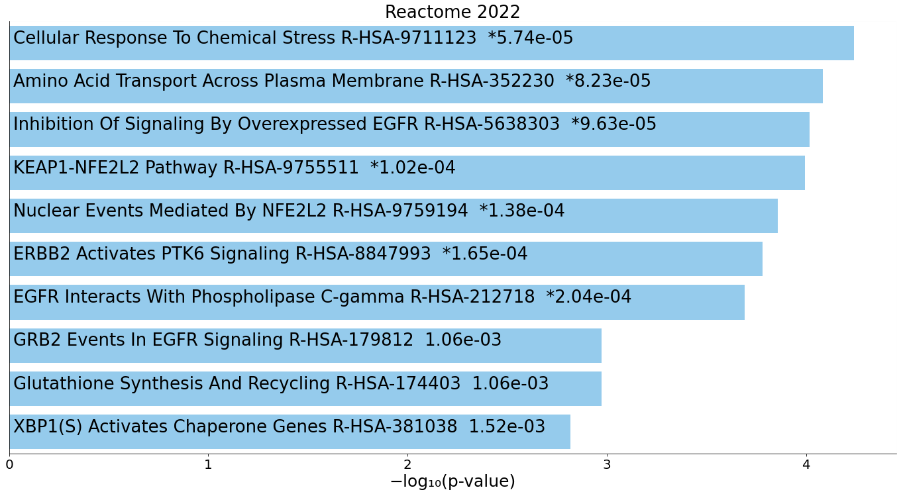
**S1B) Pathways downregulated in microstructures obtained from high-risk women’s breast tissue and exposed to LicA ex vivo.**

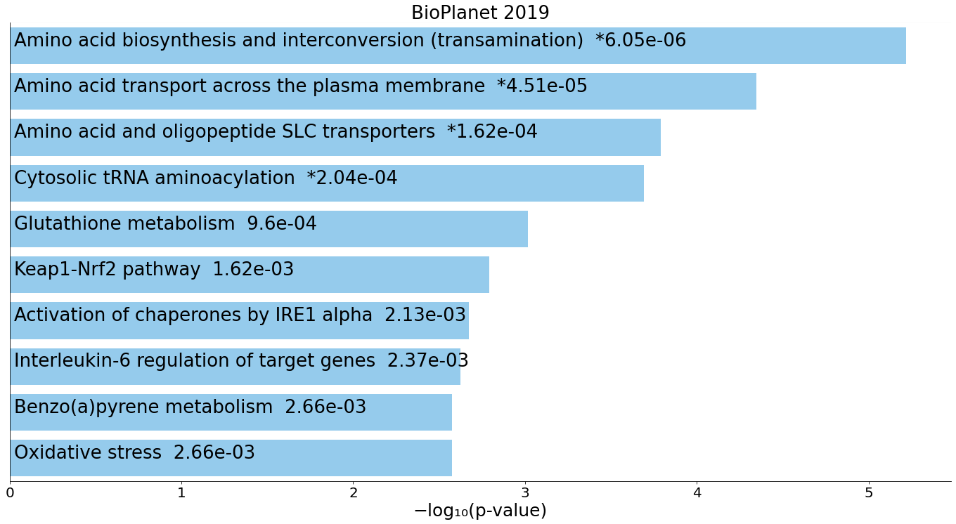

**S2) Flux through the cholesterol and steroid biosynthesis as a function of LicA exposure in microstructures obtained from high-risk women’s breast.** Microstructures from 6 subjects were treated with LicA (5 µM) for 24 h. RNA sequencing, differential gene expression, and metabolism flux analysis were performed. Orange arrows represent the direction of flux in the treated samples.

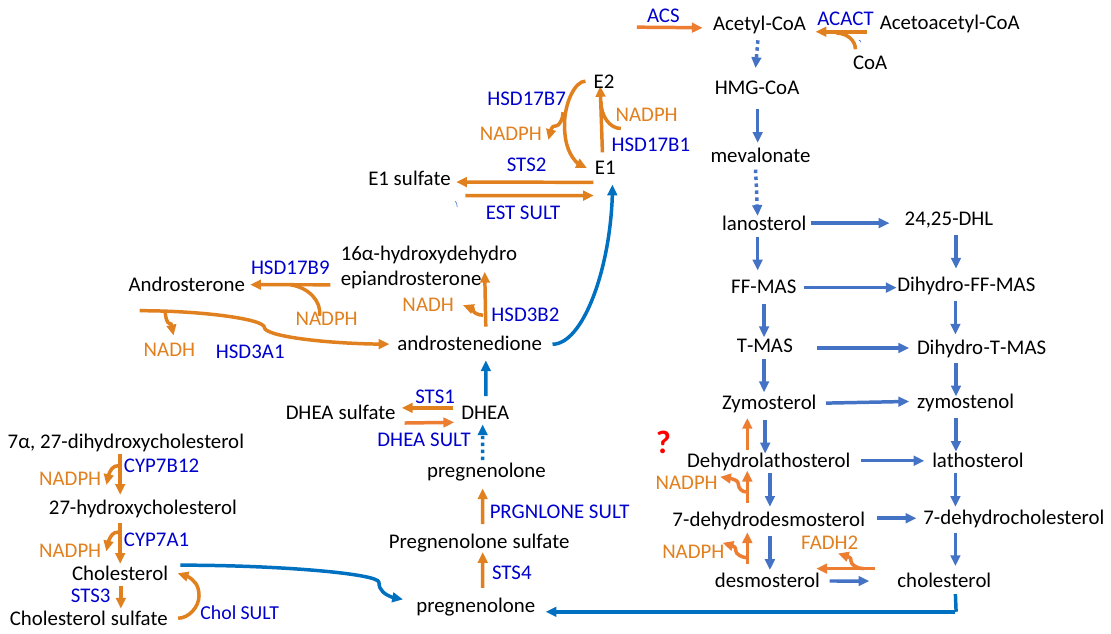

**S3A) Heatmap plot of all the identified proteins in MDA-MB-231 cells treated with DMSO in comparison to the cells treated with LicA.** MDA-MB-231 cells were incubated with DMSO or LicA (10 µM) for 24 h. Cells were lysed and undergone protein denaturation using the PISA proteomics protocol. After TMT labeling LC-MS/MS was conducted to capture the stabilized and destabilized proteins.

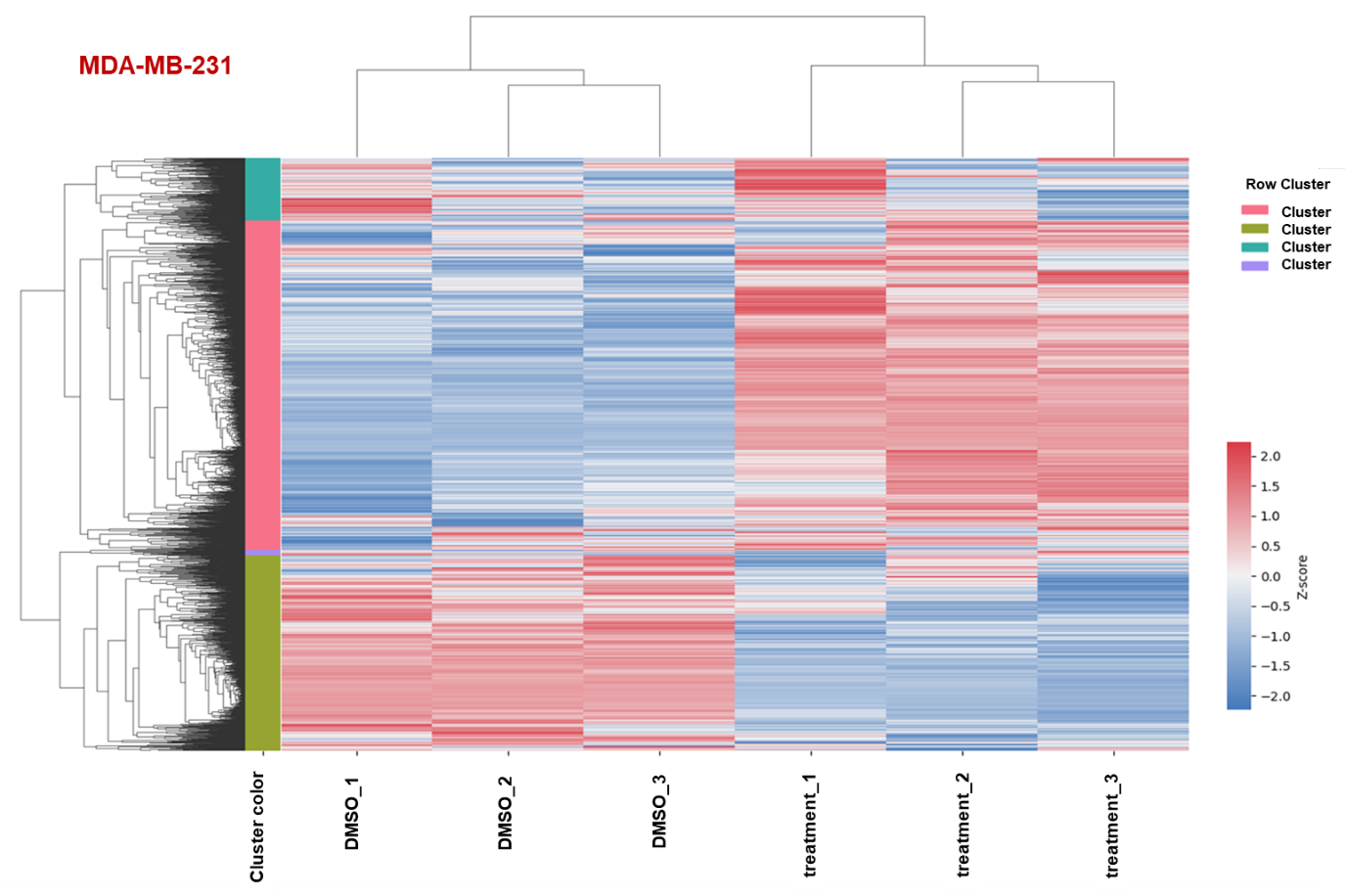

**S3B) Heatmap plot of all the identified proteins in MCF-7 cells treated with DMSO in comparison to the cells treated with LicA.** MCF-7 cells were incubated with DMSO or LicA (10 µM) for 24 h. Cells were lysed and undergone protein denaturation using the PISA proteomics protocol. After TMT labeling LC-MS/MS was conducted to capture the stabilized and destabilized proteins.

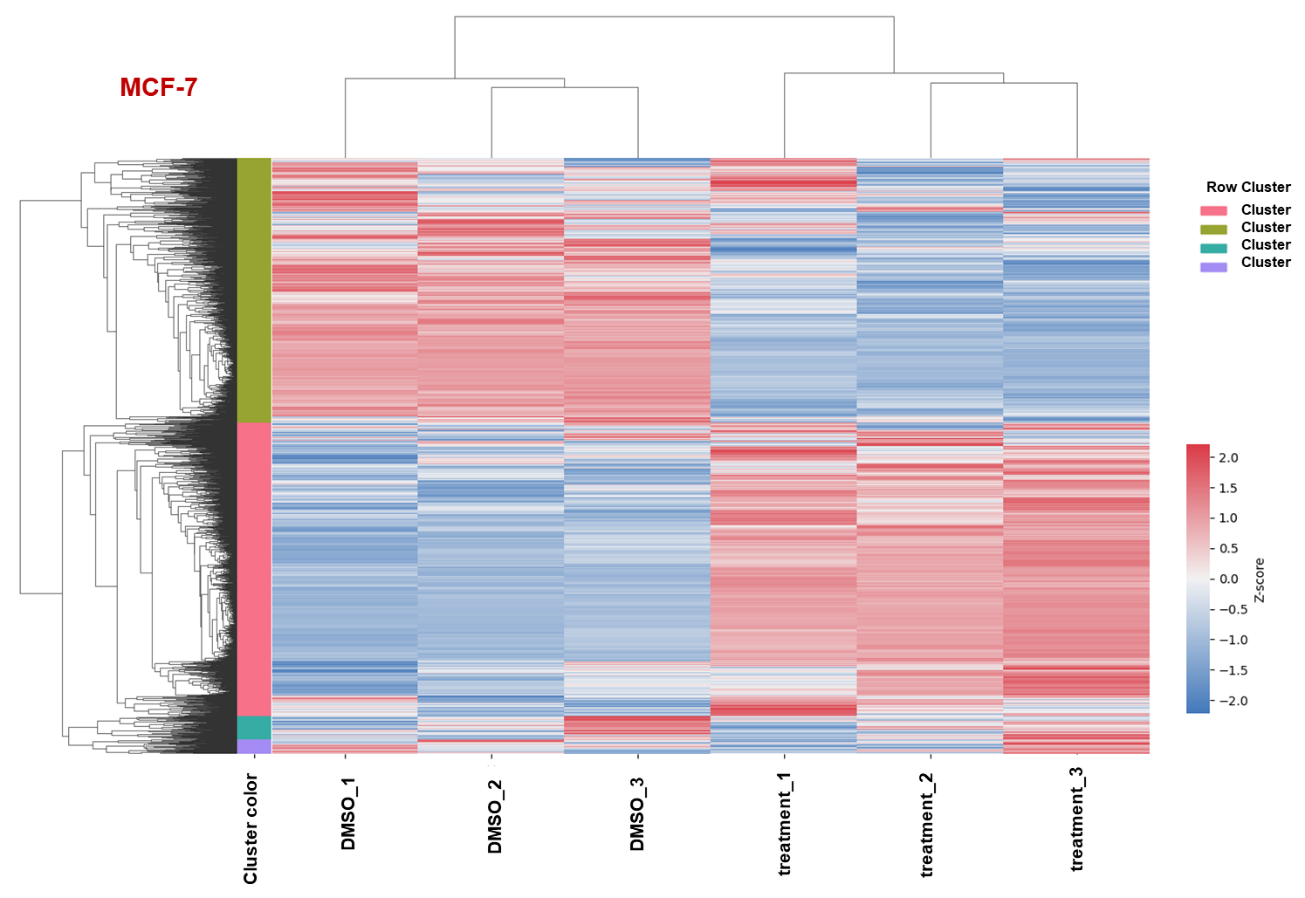
